# Can model-free reinforcement learning operate over information stored in working-memory?

**DOI:** 10.1101/107698

**Authors:** Carolina Feher da Silva, Yuan-Wei Yao, Todd A. Hare

## Abstract

Model-free learning creates stimulus-response associations. But what constitutes a stimulus? Are there limits to types of stimuli a model-free or habitual system can operate over? Most experiments on reward learning in humans and animals have used discrete sensory stimuli, but there is no algorithmic reason that model-free learning should be restricted to external stimuli, and recent theories have suggested that model-free processes may operate over highly abstract concepts and goals. Our study aimed to determine whether model-free learning processes can operate over environmental states defined by information held in working memory. Specifically, we tested whether or not humans can learn explicit temporal patterns of individually uninformative cues in a model-free manner. We compared the data from human participants in a reward learning paradigm using (1) a simultaneous symbol presentation condition or (2) a sequential symbol presentation condition, wherein the same visual stimuli were presented simultaneously or as a temporal sequence that required working memory. We found a significant effect of reward on human behavior in the sequential presentation condition, indicating that model-free learning can operate on information stored in working memory. Further analyses, however, revealed that the behavior of the participants contradicts the basic assumptions of our hypotheses, and it is possible that the observed effect of reward was generated by model-based rather than model-free learning. Thus it is not possible to draw any conclusions from out study regarding model-free learning of temporal sequences held in working memory. We conclude instead that careful thought should be given about how to best explain two-stage tasks to participants.

## 1 Introduction

Reinforcement learning theory and the computational algorithms associated with it have been extremely influential in the behavioral, biological, and computer sciences. Reinforcement learning theory describes how an agent learns by interacting with its environment [1]. In a typical reinforcement learning paradigm, the agent selects an action and the environment responds by presenting rewards and taking the agent to the next situation, or state. A reinforcement learning algorithm determines how the agent changes its action selection strategy as a result of experience, with the goal of maximizing future rewards. Depending on how algorithms accomplish this goal, they are classified as model-free or model-based [1]. Model-based algorithms acquire beliefs about how the environment generates outcomes in response to their actions and select actions according to their predicted consequences. By contrast, model-free algorithms generate a propensity to perform, in each state of the world, actions that were more rewarding in previous visits to that environmental state. Model-free reinforcement learning algorithms are of considerable interest to behavioral and biological scientists, in part because they offer a compelling account of the phasic activity of dopamine neurons, but also more generally can explain many observed patterns of behavior in human and non-human animals [2, 3, 4, 5, 6, 7].

A key concept in reinforcement learning theory is the environmental state. Typically, empirical tests of reinforcement learning algorithms use discrete sensory stimuli to define environmental states. However, there is no theoretical or algorithmic constraint to define the states of the environment exclusively by sensory stimuli. State definitions may also include the agent’s internal stimuli, such as its memory of past events, thirst or hunger level, or even subjective characteristics such as happiness or sadness [1]. Thus, model-free reinforcement learning might operate over a wide variety of both external and internal factors.

Indeed, recent work suggests that model-free learning algorithms can support a large set of cognitive processes and behaviors beyond the formation of habitual response associations with discrete sensory stimuli [8, 9, 10]. For instance, it has been proposed that the model-free system can perform the action of selecting a goal for goal-directed planning [11] or conversely that a model-based decision can trigger a habitual action sequence [12, 13, 14, 15]. Model-free algorithms have also been suggested to gate working memory [16]. However, many of these important theoretical proposals about model-free algorithms have not been directly tested empirically.

Here, we determine the ability of model-free reinforcement learning algorithms to operate over states defined by information held in working memory, an internal state. Specifically, we use an experimental paradigm and computational modeling framework designed to dissociate model-free from model-based influences on behavior [17] to test if temporally separated sequences of individually uninformative cues can drive model-free learning and behavior. If an agent can store the elements of a temporal sequence in its memory to form a unique and predictive cue and use the memorized information as the state definition, then, theoretically, it can use model-free algorithms to learn the associations between a specific sequence of *individually uninformative cues* and action outcomes [18].

Our approach has several important facets. First, we use an experimental paradigm that allows us to determine not only if our participants learn from information in working memory, but also whether that learning is supported by model-based or model-free algorithms. Second, the cues in our temporal sequences are individually uninformative; in other words, any single cue in isolation provides no information about which response is correct. It is well-known that model-free algorithms can shift response associations to the earliest occurring predictor of the correct response in a temporal sequence of informative cues and can integrate predictive information across individual cues. Neither of these mechanisms is possible in our paradigm because the individual cues themselves contain no information about the previous or subsequent cues or which response is best.

Temporal pattern learning is a fundamental and early developing human cognitive ability. It allows people to form predictions about what will happen from what has happened and select their actions accordingly. Humans can learn patterns both explicitly and implicitly in the absence of specific instructions or conscious awareness [19]. Moreover, they can do so as early as two months of age [20]. In fact, people identify patterns even when, in reality, no pattern exists [21]. These empirical results together with the theoretical potential for model-free learning to operate over internal stimuli suggest that temporal pattern learning could be supported by model-free processes. However, to date, studies of reinforcement learning and decision making have focused primarily on tasks in which the relevant stimuli are presented simultaneously just prior to or at the time of decision-making, or on implicit motor sequence learning, wherein participants learn a sequence of movements automatically, without full awareness (for instance, 22, 23, 24, 25, 26). Thus, the degree to which model-free processes do in fact operate over temporal sequences or any other information stored in working memory has not yet been directly tested and compared with model-free learning from traditionally employed external, static environmental cues.

Here, we directly test whether model-free processes can access and learn from information stored in working memory. We adapted a decision-making paradigm originally developed by Daw et al. [17] that can behaviorally dissociate the influence of model-free and model-based learning on choice. The task was performed by two groups of human participants either in a simultaneous condition (i.e. static and external), wherein visual stimuli were presented simultaneously, or in a sequential condition, wherein the same visual stimuli were presented as a temporal sequence that required working memory processing.

## 2 Results

### 2.1 Determining model-free and model-based influences on choice behavior

Forty-one young adult human participants completed a behavioral task adapted from Daw et al. [17]. In our task, participants began each trial in a randomly selected initial state represented by one of four possible sequences of two symbols: AA, AB, BA, or BB (Figure 1). At this initial state, participants chose one of two possible actions: going left or going right. They were then taken to one of two possible final states, the blue state or the pink state. If they had gone left, they were taken with 0.8 probability to the final state given by the rule AA → blue, AB → pink, BA → pink, BB → blue or with 0.2 probability to the other final state. If they had gone right, they were taken with 0.8 probability to the final state *not* given by the previous rule or with 0.2 probability to the other final state. The common (most probable) transitions between the initial and final states are shown in Figure 2. To predict the final state accurately, participants had to know both elements of the sequence. If they knew only one, the final state might have been either blue or pink with 0.5 probability and they would not be able to perform above chance. This feature is key and separates our work from others in which each element of a sequence is predictive on its own.

**Figure 1:**
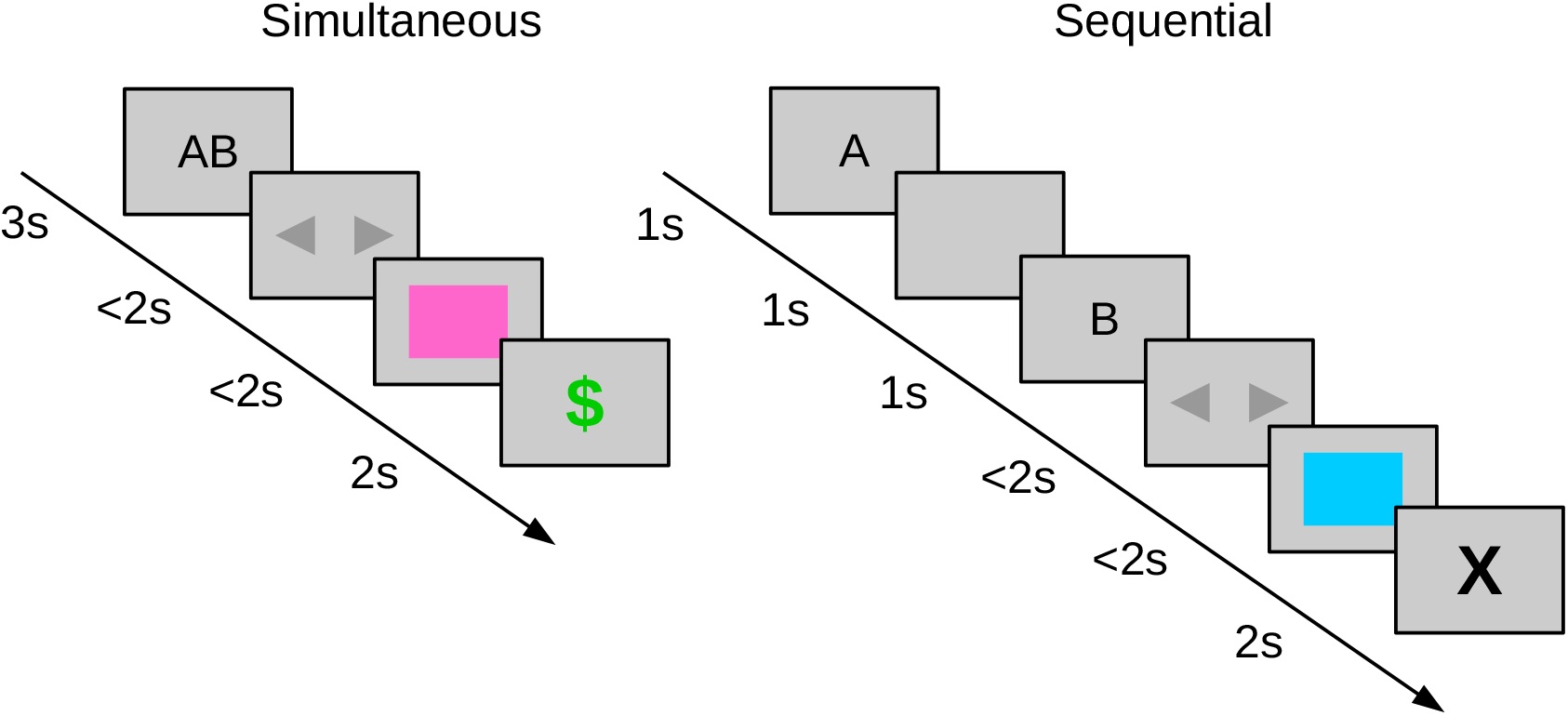
Timelines of events in a trial. The two symbols that represent the initial state are presented simultaneously in the simultaneous condition (left) and separately as a temporal sequence in the sequential condition (right). In this example, AB is the initial state. The simultaneous condition participant goes to the pink final state and receives a reward (signaled by the green $ symbol). The sequential condition participant goes to the blue final state and does not receive a reward (signaled by the black X symbol).

**Figure 2:**
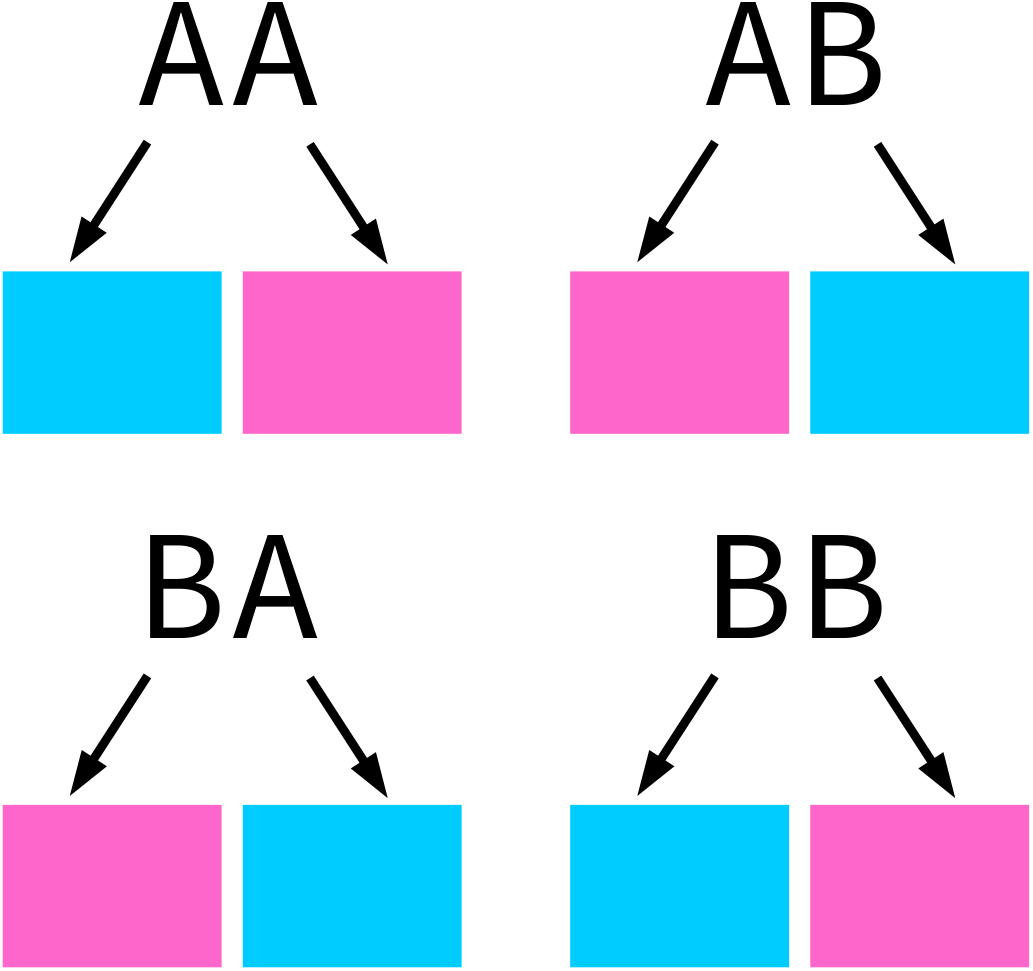
Common state transitions in the behavioral task’s model. These graphics highlight the uninformative nature of each single element (i.e. A or B symbols) in the simultaneous or sequential cues. Knowledge of only the first or final element of the combined cue provides no indication of how likely the right and left responses are to lead to a specific state.

One of the final states delivered a monetary reward with 0.7 probability and the other with 0.3 probability. The optimal strategy was to always select the action that led with 0.8 probability to the final state with 0.7 reward probability. Initially, participants were instructed to learn the common transitions between the initial and final states in the absence of rewards. They were told that each final state might be rewarded with different probabilities, but not what the probabilities were nor that they were fixed. The task comprised 250 trials and participants received the total reward they obtained at the end.

Twenty-one participants were randomly allocated to a simultaneous condition and twenty to a sequential condition (Figure 1). In the simultaneous condition, both symbols that represented the initial state were displayed simultaneously on the screen. In the sequential condition, each symbol was displayed consecutively by itself, as a temporal sequence. The specific objective of this study was to determine if participants in the sequential condition could use states represented in working memory to learn the task in a model-free way or if their learning was necessarily model-based. The simultaneous condition is already known to support model-free learning as well as model-based learning [17, 27, 28, 29, 30]. We thus sought to determine the difference between the standard simultaneous and working-memory dependent sequential conditions.

The two-stage task we used can differentiate between model-free and model-based learning because algorithms that implement them make different predictions about how a reward received in a trial impacts a participant’s choices in subsequent trials. The SARSA (*λ* =1) model-free algorithm learns this task by strengthening or weakening associations between initial states and initial-state actions depending on whether the action is followed by a reward or not [1]. Therefore, it simply predicts that an initial-state action that resulted in a reward is more likely to be repeated in the next trial with the same initial state [17]. On the other hand, the model-based algorithm considered in this study uses an internal model of the task’s structure to determine the initial-state choice that will most likely result in a reward [17]. To this end, it considers which final state, pink or blue, was most frequently rewarded in recent trials and selects the initial-state action, left or right, that will most likely lead there. Therefore, the model-free algorithm predicts that the participant will choose the mostly frequently rewarded *action* in past trials with the same initial state, while the model-based algorithm predicts that the participant will choose the action with the highest probability of leading to the mostly frequently rewarded *final state* in past trials, regardless of their initial states.

The model-free and model-based algorithms thus generate different predictions about the *stay probability*, which is the probability that in two consecutive trials the participant will stay with their first choice and take the same initial-state action in the second trial. For instance, if the participant chose left in two consecutive trials, this was considered a stay. The model-free and model-based predictions are different if the letters presented in one trial are the same or different than the letters presented in the other trial, so we ran four separate analyses on the data from each condition, dividing consecutive trial pairs into four subsets: “same letters” if both letters presented in the first trial are the same as the letters presented in the second trial (for example, AB for the first trial and AB again for the second trial), “same first letter” if the first letters presented in each trial are the same but the second letters are different (for example, AB and AA), “same second letter” if the second letters are the same but the first letters are different (for example, AB and BB), and “different letters” if both the first letters and the second letters are different (for example, AB and BA).

In all cases, we analyzed the data using Bayesian hierarchical logistic regression analyses. In addition to examining the stay choice probabilities, we directly examined the logistic regression coefficients for each condition and trial pair subset. Because in our task the mean reward probability associated with one final state is higher than the mean reward probability associated with the other final state, we did not use the logistic regression model proposed by Daw et al. [17]—as several studies demonstrate, if the reward probabilities are not the same, a reward by transition interaction does not uniquely characterizes model-based agents, but also appears in purely model-free results [31, 32, 33, 34]. We thus used instead an extended logistic regression model we had previously proposed that corrects for different reward probabilities by adding two control predictors: a binary variable that indicates whether or not the chosen initial-state action in the first trial leads commonly to the final state with the highest reward probability, and a binary variable that indicates whether or not the agent visited in the first trial the final state with the highest reward probability [34]. For comparison with the behavior of human participants, we fitted model-free and model-based algorithms to the experimental data, used the obtained parameter estimates to simulate purely model-free and purely model-based agents performing our task, and analyzed the resulting data using the same logistic regression procedure. The stay probabilities obtained for both simulated agents and human participants are shown in Figures 3, 4, 5, and 6, and the logistic regression coefficients obtained for both simulated agents and human participants are shown in Figures 7, 8, 9, and 10.

**Figure 3:**
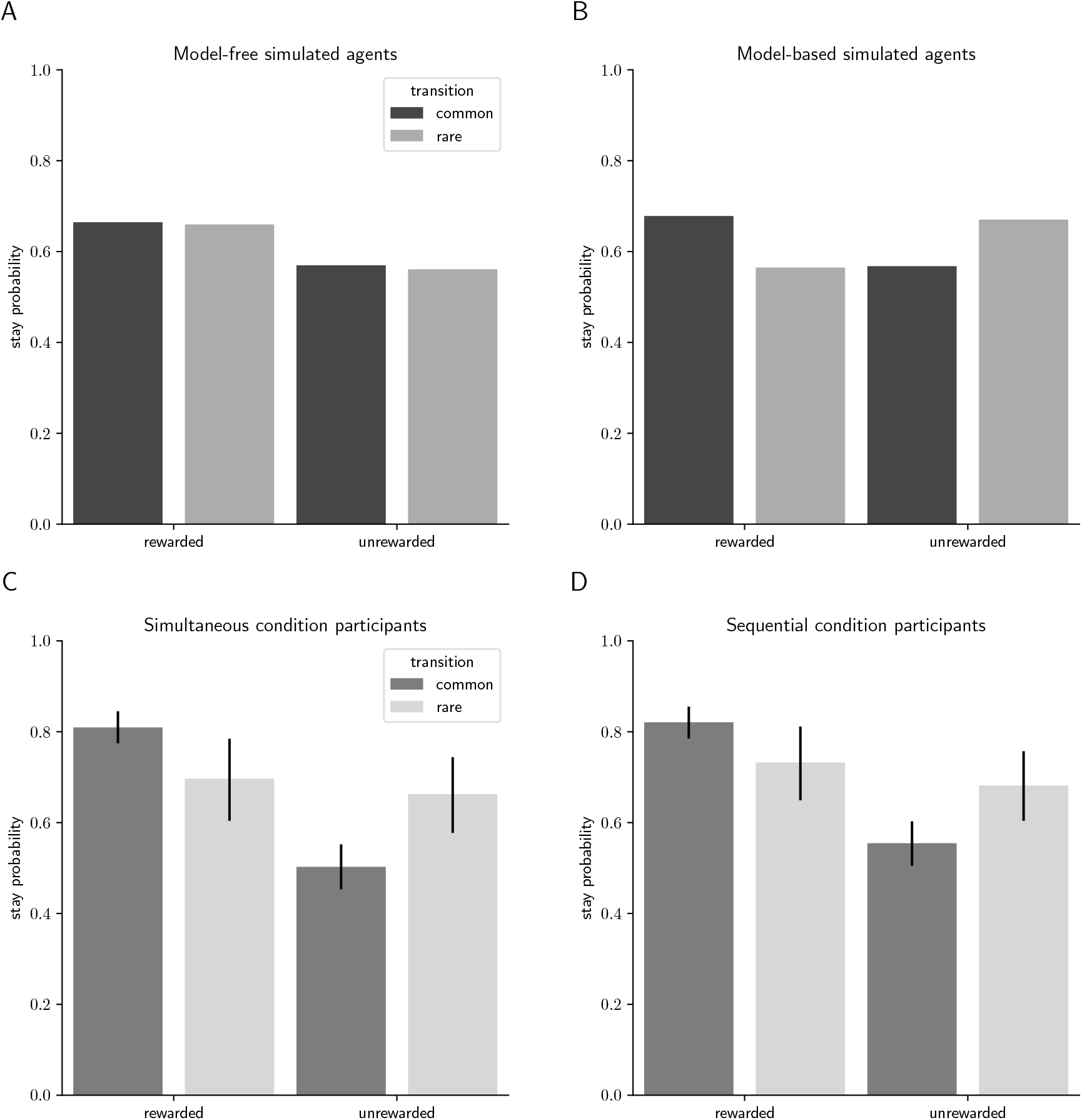
Stay probabilities of simulated agents and human participants for consecutive trial pairs in the “same letters” subset. A- Stay probabilities of purely model-free simulated agents. B- Stay probabilities of purely model-based simulated agents. C- Stay probabilities of human participants in the simultaneous condition. D- Stay probabilities of human participants in the sequential condition. The error bars correspond to the 95% credible interval.

**Figure 4:**
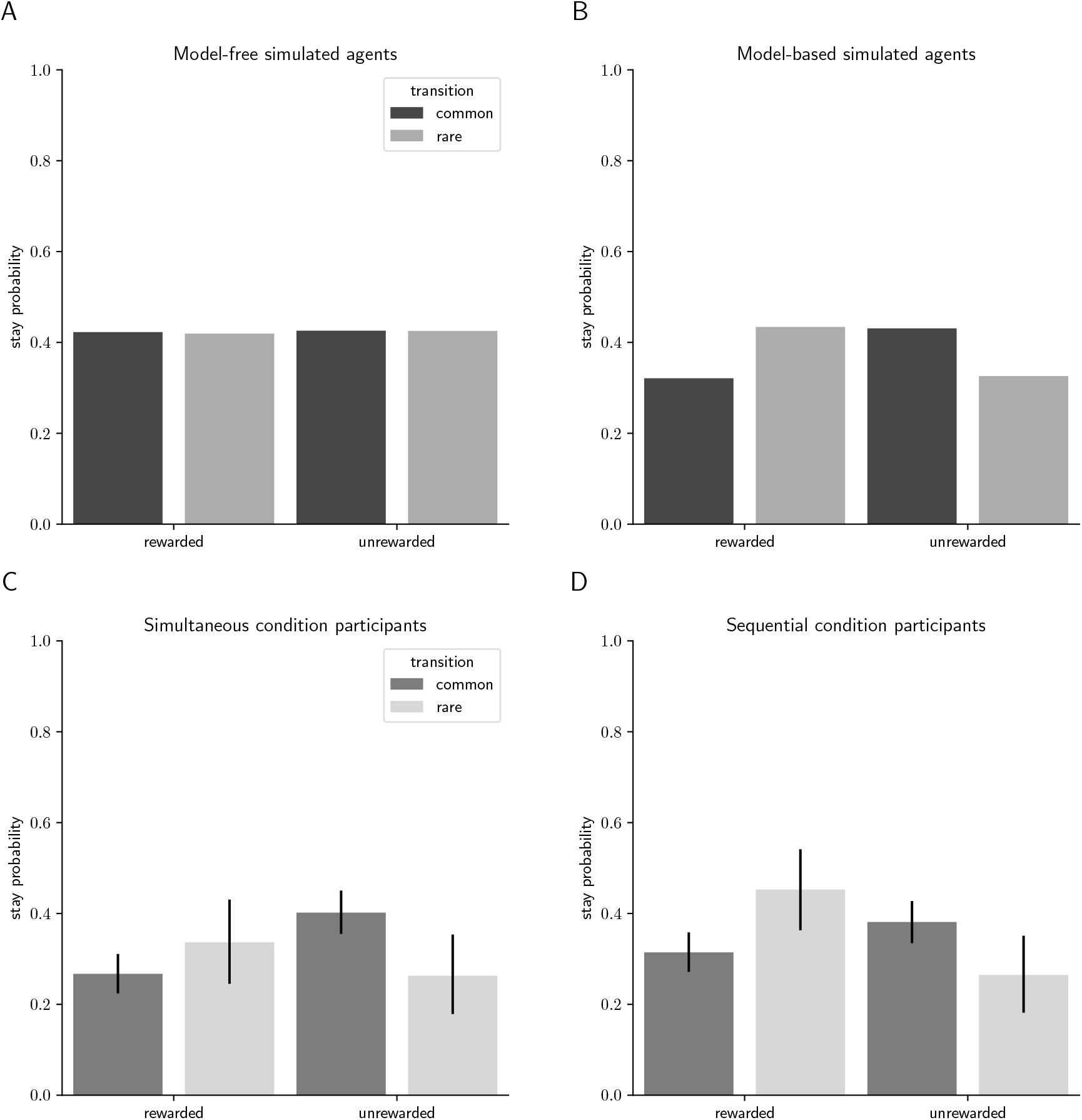
Stay probabilities of simulated agents and human participants for consecutive trial pairs in the “same first letter” subset. A- Stay probabilities of purely model-free simulated agents. B- Stay probabilities of purely model-based simulated agents. C- Stay probabilities of human participants in the simultaneous condition. D- Stay probabilities of human participants in the sequential condition. The error bars correspond to the 95% credible interval.

**Figure 5:**
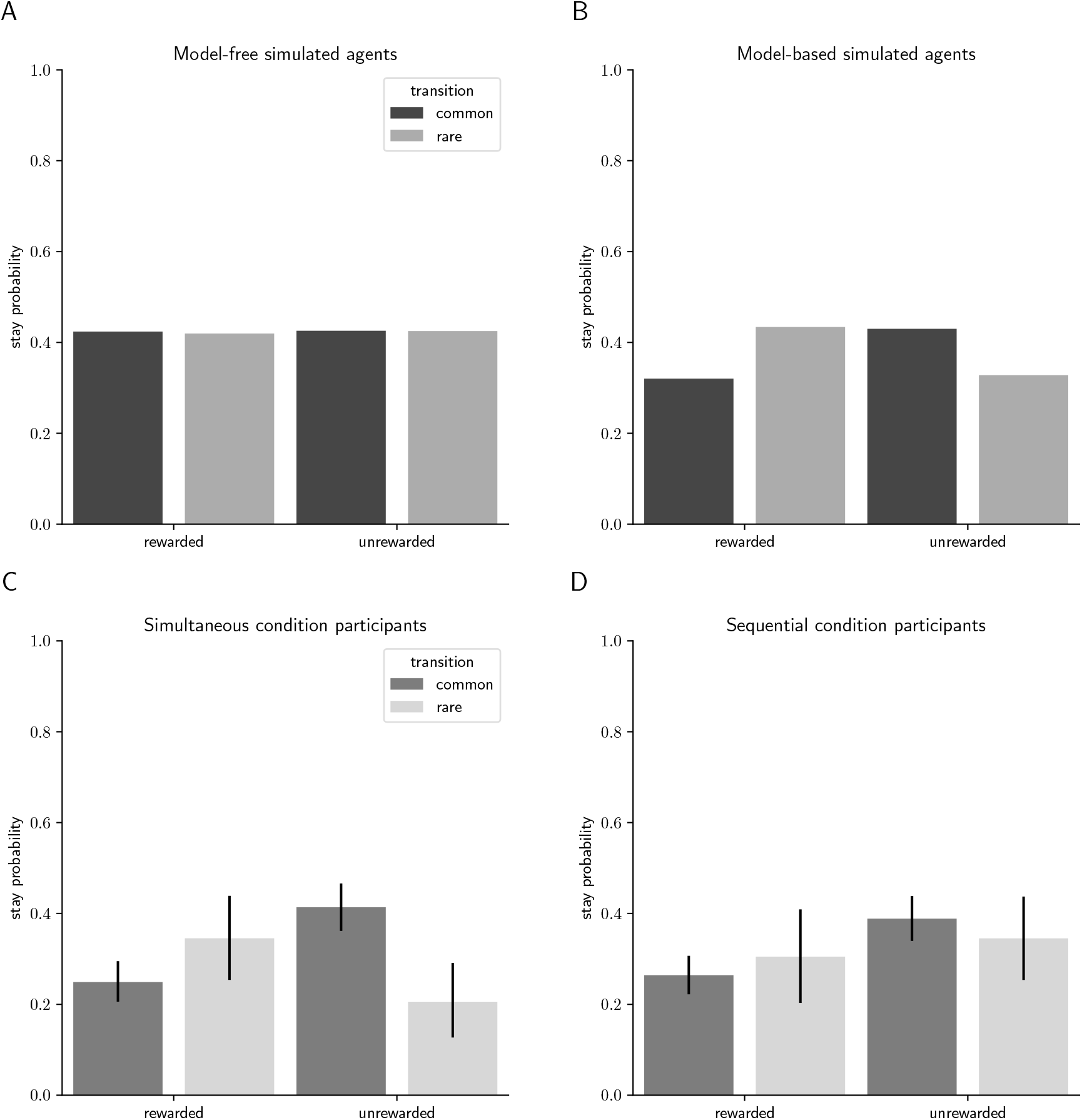
Stay probabilities of simulated agents and human participants for consecutive trial pairs in the “same second letter” subset. A- Stay probabilities of purely model-free simulated agents. B- Stay probabilities of purely model-based simulated agents. C- Stay probabilities of human participants in the simultaneous condition. D- Stay probabilities of human participants in the sequential condition. The error bars correspond to the 95% credible interval.

**Figure 6:**
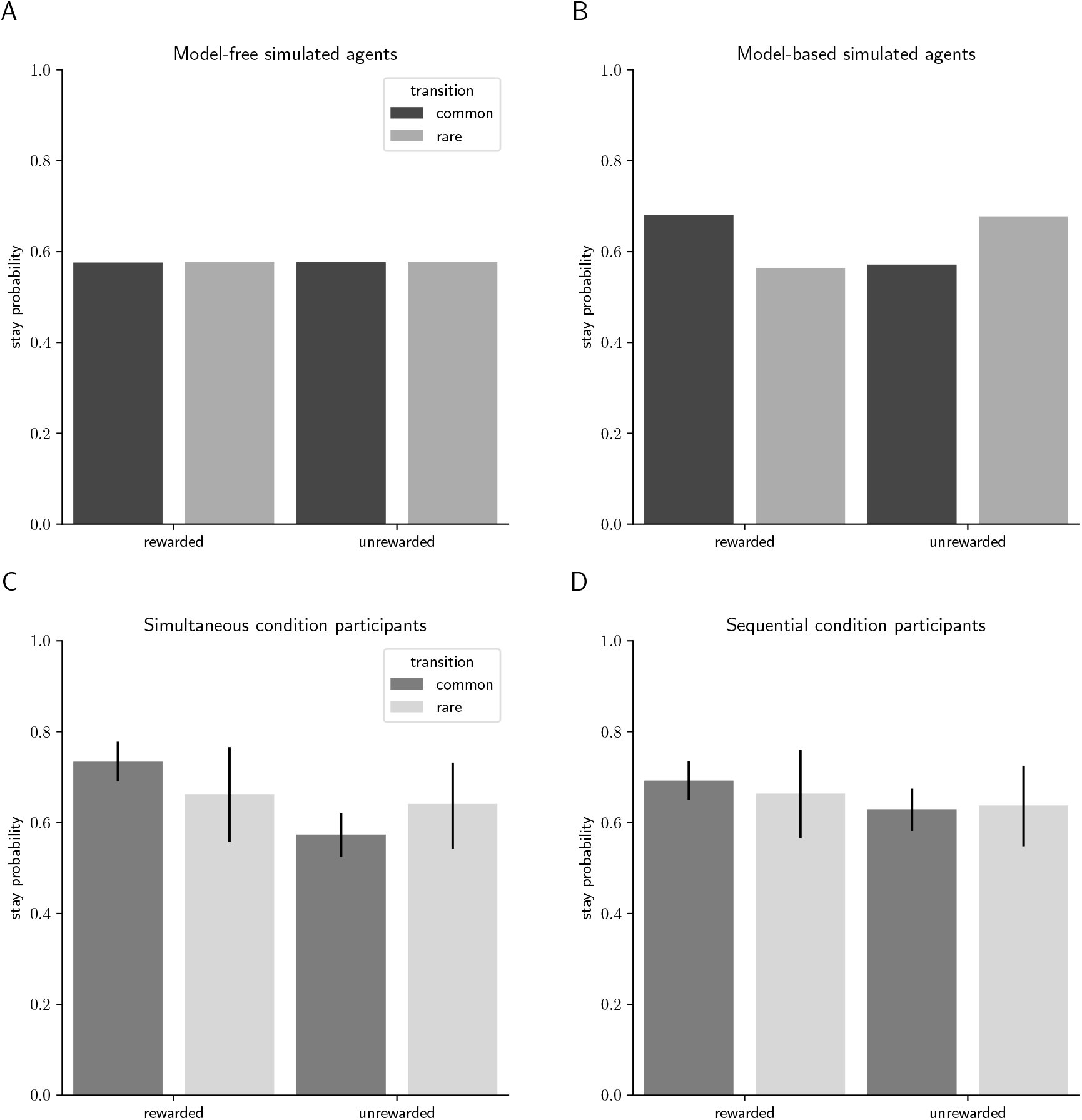
Stay probabilities of simulated agents and human participants for consecutive trial pairs in the “same letters” subset. A- Stay probabilities of purely model-free simulated agents. B- Stay probabilities of purely model-based simulated agents. C- Stay probabilities of human participants in the simultaneous condition. D- Stay probabilities of human participants in the sequential condition. The error bars correspond to the 95% credible interval.

**Figure 7:**
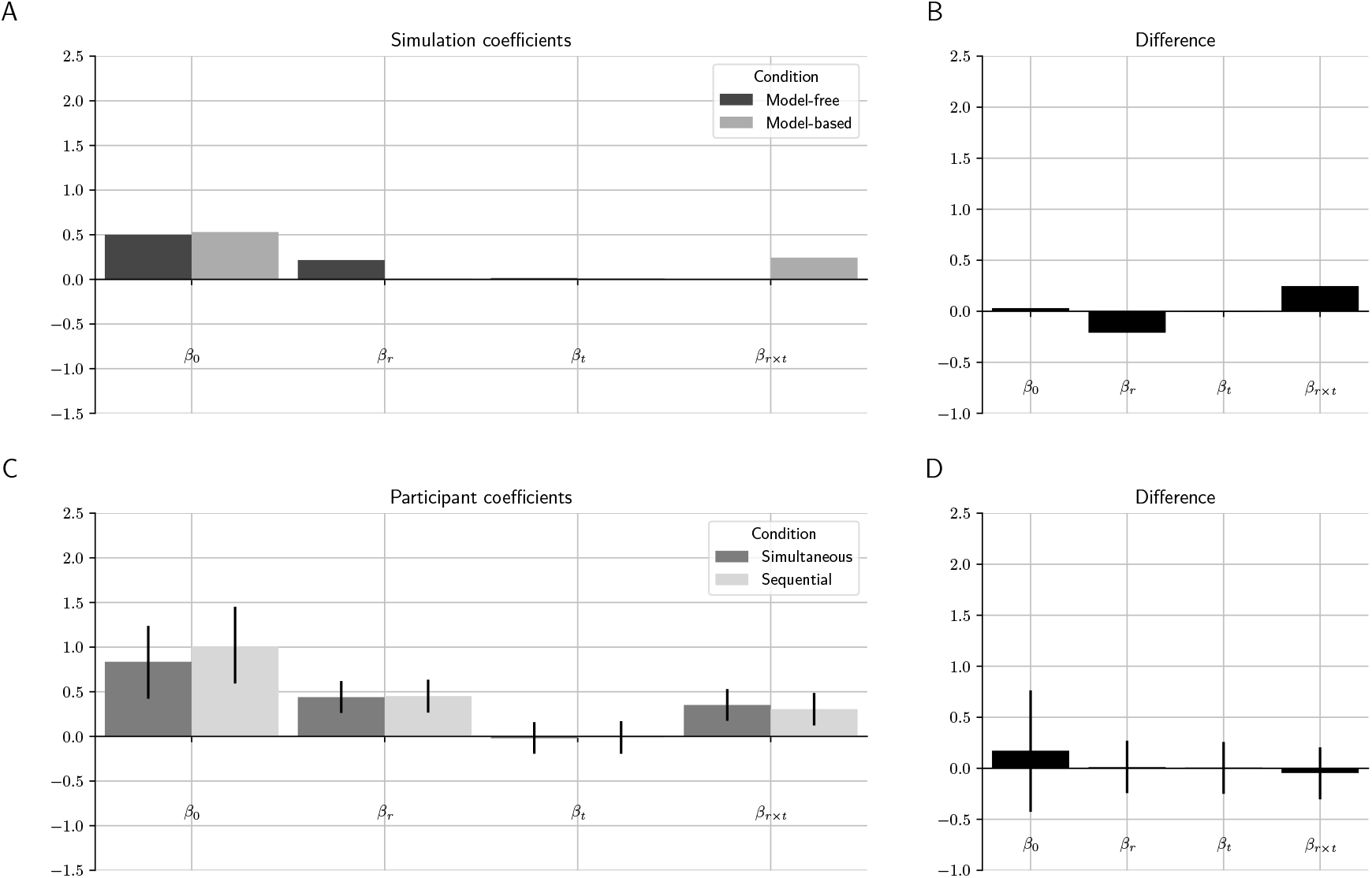
Logistic regression coefficients of simulated agents and human participants for consecutive trial pairs in the “same letters” subset. A- Logistic regression coefficients of purely model-free and purely model-based simulated agents. B- Difference between the coefficients of purely model-based and purely model-free simulated agents. C- Logistic regression coefficients of human participants in the simultaneous and sequential conditions. D- Difference between the coefficients of human participants in the simultaneous and sequential conditions. The error bars correspond to the 95% credible interval.

**Figure 8:**
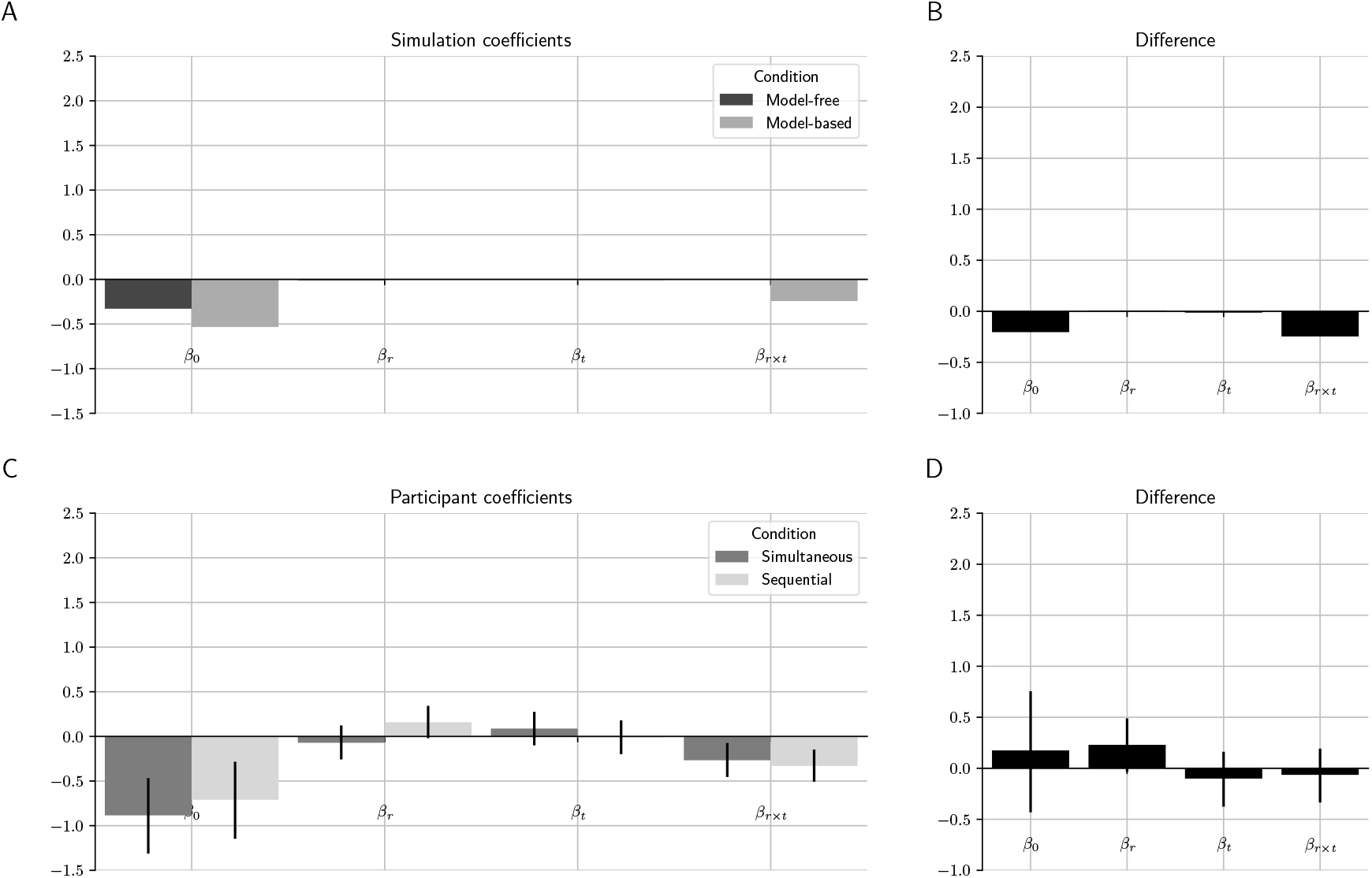
Logistic regression coefficients of simulated agents and human participants for consecutive trial pairs in the “same first letter” subset. A- Logistic regression coefficients of purely model-free and purely model-based simulated agents. B- Difference between the coefficients of purely model-based and purely model-free simulated agents. C- Logistic regression coefficients of human participants in the simultaneous and sequential conditions. D- Difference between the coefficients of human participants in the simultaneous and sequential conditions. The error bars correspond to the 95% credible interval.

**Figure 9:**
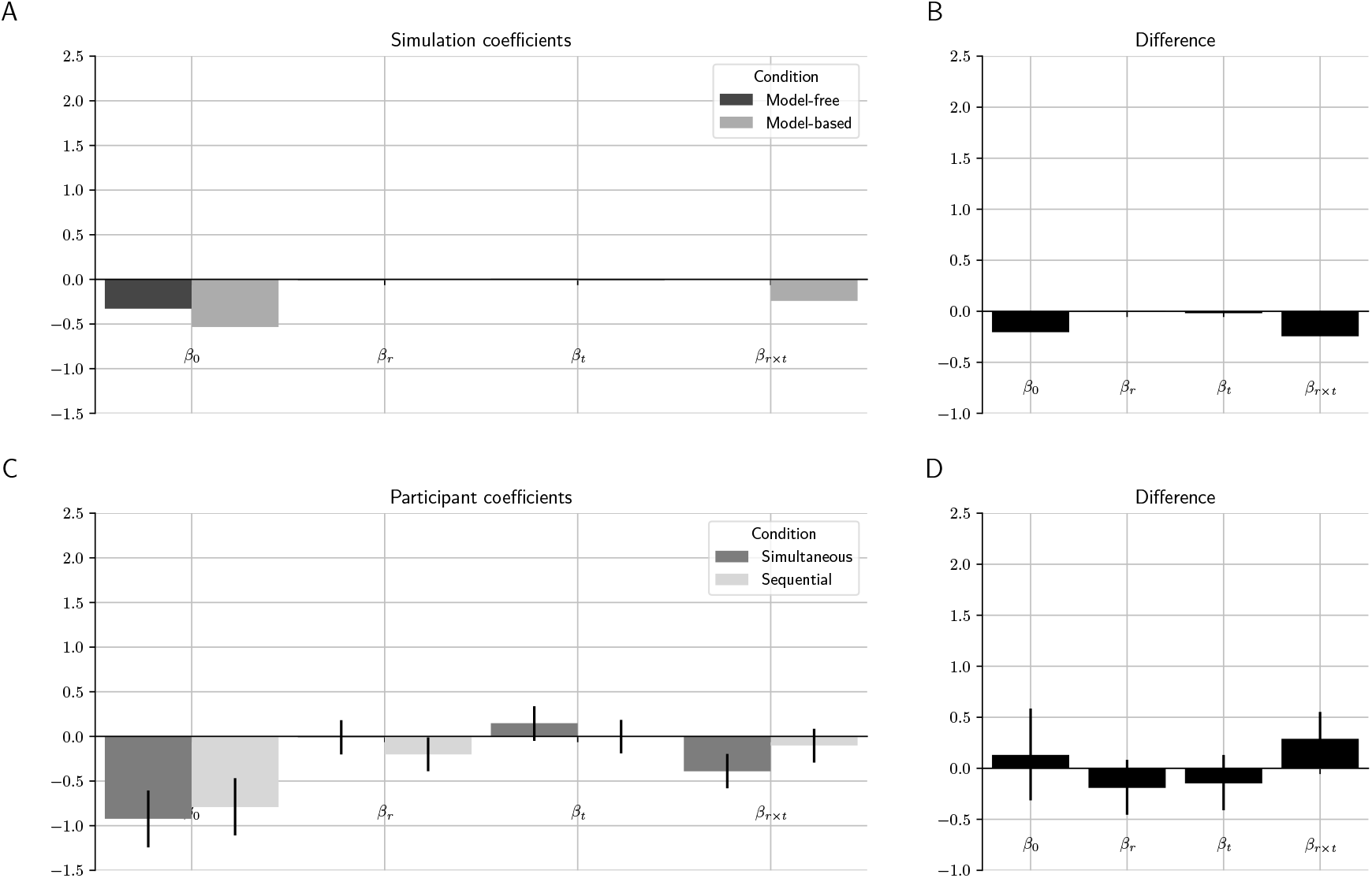
Logistic regression coefficients of simulated agents and human participants for consecutive trial pairs in the “same second letter” subset. A- Logistic regression coefficients of purely model-free and purely model-based simulated agents. B- Difference between the coefficients of purely model-based and purely model-free simulated agents. C- Logistic regression coefficients of human participants in the simultaneous and sequential conditions. D- Difference between the coefficients of human participants in the simultaneous and sequential conditions. The error bars correspond to the 95% credible interval.

**Figure 10:**
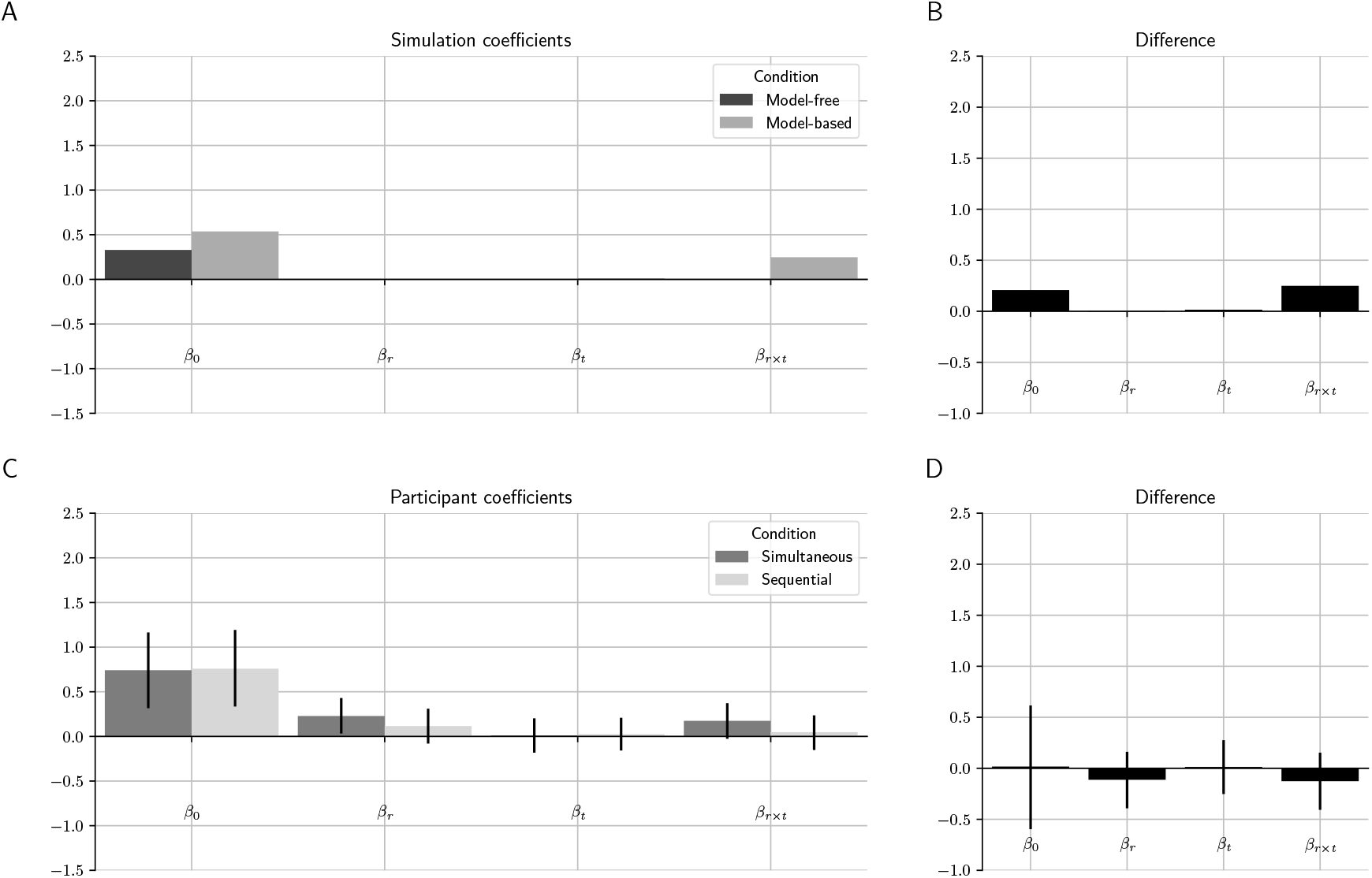
Logistic regression coefficients of simulated agents and human participants for consecutive trial pairs in the “different letters” subset. A- Logistic regression coefficients of purely model-free and purely model-based simulated agents. B- Difference between the coefficients of purely model-based and purely model-free simulated agents. C- Logistic regression coefficients of human participants in the simultaneous and sequential conditions. D- Difference between the coefficients of human participants in the simultaneous and sequential conditions. The error bars correspond to the 95% credible interval.

As can be seen in Figure 7A, if the letters are the same, for example AB for both trials, the model-free prediction is that the stay probability will increase if the first trial was rewarded and decrease if it was not; i.e., model-free learning creates a positive reward effect. The model-based prediction, on the other hand, is that the stay probability will increase if either the first trial was rewarded and the transition was common or the first trial was unrewarded and the transition was rare, and decrease otherwise; i.e. model-based learning creates a positive reward by transition interaction [17]. If the consecutive trials have different initial-state letters, the predictions will be different depending on the condition (simultaneous or sequential) and the assumed hypothesis regarding model-free learning of temporal sequences. In the simultaneous condition, the model-free prediction is that the stay probability will not change, because learning does not generalize among different initial states (Figures 8A, 9A, and 10A). In the sequential condition, if we assume that model-free learning *can* learn from temporal sequences, then the prediction is that the stay probability will also not change (Figures 8A, 9A, and 10A). If, however, we assume that model-free learning *cannot* learn from temporal sequences, then the model-free system may associate the first letter or the second letter to previously received rewards. Assuming, for example, that the second letter is associated with rewards, if the two consecutive trials have the same second letter, the stay probability should increase if the previous trial was rewarded and decrease if the previous trial was unrewarded; if the two consecutive trials have different second letters, however, the stay probability should not change. The simulated results for the latter hypothesis are not shown; in the Figures above, we assumed that model-free learning *can* learn from temporal sequences. For model-based learning, the prediction is that the reward by transition interaction will the positive if both letters are the same or both letters are different for the two trials (for example, the first trial’s letters are AB and the second trial’s letters are either AB or BA—see Figures 7A and 10A), because in this case the common and rare transitions are the same for both trials. If one letter is the same but the other letter is different (for example, the first trial’s letters are AB and the second trial’s letters are AA or BB—see Figures 8A and 9A), the model-based prediction is that the reward by transition interaction will be negative, because in this case the common and rare transitions are switched between the trials, so if the left action commonly leads to pink state in the first trial, for instance, it commonly leads to the blue state in the second trial.

For our sample of the human participants and trial pairs in the “same letters” subset, behavior was influenced by both reward and reward by transition interaction regardless of whether the states were defined by external sensory cues or internal working-memory representations (Figures 7C). We thus found no evidence that sequentially presented, working-memory-dependent state cues shift the balance of model-based and model-free effects on choice behavior compared to traditional, static, external cues. However, the results obtained for other trial pair subsets show unpredicted effects, namely: (1) there is a negative effect of reward for the sequential condition in the “same second letter” subset (Figure 9C); the estimated value of this coefficient is −0.20 (95% CI [−0.39, −0.01]); and (2) there is a positive effect of reward for the simultaneous condition in the “different letters” subset (Figure 10C); the estimated value of this coefficient is 0.23 (95% CI [0.02, 0.43]). Because of these unexpected results, we decided to replicate our experiment using a task that had geometric figures rather than letters to identify the different initial states (see Appendix on page 29). 32 human participants performed that task in both the simultaneous and sequential conditions. We again observed in the replicated data a negative reward effect for the sequential condition in the “same first letter” and “same second letter” subsets, as well as a positive reward effect for both the sequential and the simultaneous condition in the “different letters” subset.

## 3 Discussion

In this study, we empirically tested the hypothesis that human participants can develop model-free associations between temporal sequences of stimuli stored in working memory and a motor response. To that end, we developed a behavioral task based on a previous decision-making paradigm that can determine the model-free and model-based influences on choice [17]. The participants in the simultaneous condition performed this task with the two visual symbols presented together simultaneously and those in the sequential condition performed it with the same two visual symbols presented as a temporal sequence that had to be held in working memory. A key element of our experimental paradigm is that the individual symbols within each temporal sequence convey no information about the best response in isolation. This fact rules out the possibility that the sequential condition’s model-free effect is due to an association between a single symbol in the sequence and a response rather than one between the entire sequence and a response. Each sequence element is completely uninformative by itself: it cannot predict reward delivery above chance. Therefore, the task cannot be learned by simple stimulus-response associations with individual symbols in the temporal sequence.

At first glance, our results support the hypothesis that model-free learning can operate on stimuli stored in working memory. Two findings, however, cannot be explained by the assumed model of hybrid reinforcement learning, adapted to the two-stage task by Daw et al. [17]. Since model-free learning is assumed to be unable to generalize between distinct states (see Doll et al. 35, Kool et al. 36 for example studies that critically depend on this assumption) and model-based learning is assumed to generate only a reward by transition interaction, there should not be a reward effect for consecutive trials with different initial-state symbols. Yet, we observed a positive reward effect for trial pairs in the “different letters” subset both in the data presented here and in the follow-up replication study using a different initial-state representation. A possible explanation for this finding is that, after all, model-free learning is able to generalize between different state representations. It is possible that participants reduced the two-letter sequence to an abstract representation such as “the two letters were the same” (either AA or BB) or “the two letters were different” (either AB or BA). This abstraction is sufficient to determine the common and rare transitions, and we know from direct reports that at least some participants used it to memorize the transition rules. If model-free learning can operate on stimuli stored in working memory, it is also conceivable it can also operate on abstract representations stored in working memory. However, the use of abstract state representations cannot explain our second unpredicted finding: a negative effect of reward observed for the sequential condition in the “same second letter” subset and, in the replication study, also in the “same first letter” subset. Under the assumed model of hybrid learning, the reward effect can never be negative. The TD(*λ* = 1) algorithm used here to model model-free learning in the brain foresees no circumstances under which rewarding one action would *decrease* the probability of choosing that action again in the future.

The unpredicted reward effects we observed in some analyses raise a question about the predicted reward effect observed in other analyses: Does a reward effect truly indicate model-free learning in our data set? Is it not possible that at least some of these effects are generated by model-based learning instead? It is commonly assumed that model-based learning does not generate a reward effect, because it is assumed that participants make model-based decisions using a specific model of the task structure. It is possible, however, that the model they are using is different from the assumed one and can generate positive as well as negative reward effects. For example, a participant might think that their initial-state choices influence the reward probability, even if they are told this is not the case—they might have misunderstood or forgotten the instructions or thought the instructions were misleading.

Given that at least some of the observed reward effects may be generated by model-based rather than model-free learning, we cannot conclude that our data presents evidence for or against the hypothesis that model-free learning can operate over information held in working memory. In order to study this or other hypotheses involving model-free learning, it is crucial that participants are using a model of the task structure for model-based learning that does not generate reward effects. Future research may thus concentrate on developing more detailed and precise instructions, as well as tutorials and tests, to make sure that participants really understood the task and what they have to do. It is also essential that the data are checked for violations of the assumed model using multiple analyses.

## 4 Methods

### 4.1 Participants

Forty-one healthy young adults participated in the experiment, 21 (13 female) randomly assigned by a random number generator to the simultaneous condition and 20 (13 female) to the sequential condition. The inclusion criterion was speaking English and no participants were excluded from the analysis. The sample size was chosen by the precision for research planning method [37, 38], by comparing the estimated differences between participant groups in the logistic regression analysis with those between model-free and model-based simulated agents.

The experiment was conducted in accordance with the Zurich Cantonal Ethics Commission’s norms for conducting research with human participants, and all participants gave written informed consent.

### 4.2 Task

The task’s state transition model defines four possible initial states, which were randomly selected with uniform distribution in each trial and represented by four different stimuli, each composed of two symbols: AA, AB, BA, or BB. At the initial state, two actions were available to the participant: pressing the left or the right arrow keys. By pressing one of the keys, the participant was taken to a final state, which might be either the blue state or the pink state. If the left arrow key was pressed, the participant was taken to the final state given by the rule AA → blue, AB → pink, BA → pink, BB → blue with 0.8 probability or to the other state with 0.2 probability; if the right arrow key was pressed, the participant was taken to the final state not given by the previous rule with 0.8 probability or to the other state with 0.2 probability. There was no choice of action at the final state, but participants were required to make a button press to potentially earn the reward. Each final state was rewarded according to an associated probability, which was 0.7 for one state and 0.3 for the other. The highest reward probability was associated with the blue state for half of the participants and to the pink state for the other half. Participants were told that each final state might be rewarded with different probabilities, but not what the probabilities were nor that they were fixed.

In contrast with our task design, in which the final states’ reward probabilities were fixed, in the original task design proposed by Daw et al. [17] the reward probabilities slowly drifted over time, because those authors were interested in the trade-off between model-based and model-free mechanisms, which is assumed to happen on the basis of their relative uncertainties. In this study we were interested instead in testing if model-free learning of temporal patterns is possible and keeping the task environment stable helps making the model-free associations stronger and more likely to influence choice [39, 40].

Participants were initially instructed to learn the common transitions between the initial and the final states in the absence of reward. Participants then performed the task defined by the model above in the simultaneous or sequential condition. Half of the participants were randomly allocated to the simultaneous condition and the other half to the sequential condition (Figure 1). In the simultaneous condition, both symbols that define the initial state were displayed simultaneously on the screen for 3 seconds. In the sequential condition, each symbol is an element of a sequence and each element was presented for 1 second, but never conjointly, and with a 1-second delay (blank screen) in between. Two triangles pointing left and right then appeared and the participant was given 2 seconds to make a decision about whether to press the left or the right arrow keys; if they did not press any keys, the word SLOW was displayed for 1 second, and the trial was aborted and omitted from analysis. A blue or pink rectangle appeared immediately afterward, indicating the final state. The participant then pressed the up-arrow key and, if the final state was rewarded, a green dollar sign appeared on the screen for 2 seconds; otherwise, a black X appeared for 2 seconds. The task comprised 250 trials, with a break every 50 trials, and participants received the total reward they obtained by the end of the task (0.18 CHF per reward).

### 4.3 Model-free algorithm

The SARSA model-free algorithm with replacing eligibility traces [1, 17] was used to simulate model-free learning agents. For each action *a* and state *s*, it estimated the value *Q*(*s, a*) of performing that action in that state. The task’s initial states *s_i_* were AA, AB, BA, and BB, and the actions *a_i_* available at the initial states were *left* and *right*. The final states were *pink* and *blue*, and the only action *a_f_* available at those states was *up*. The initial value of *Q*(*s, a*) for every state and action was 0.5. In each trial *t*, the simulated agent at the initial state *s_i_* chose *left* as its initial-state action with probability *p_left_* and *right* with probability 1 − *p_left_*, according to the following equation:

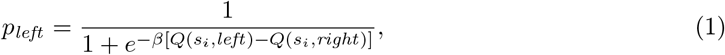

where *β* > 0 is an inverse temperature parameter that determines the algorithm’s propensity to choose the option with the highest estimated value. After the final state *s_f_* was observed and a reward *r* ∈ {0, 1} was received, state-action values were updated according to the following equations:

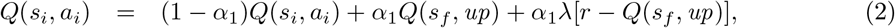

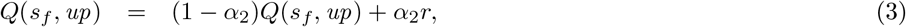

where 0 ≤ *α*_1_, *α*_2_, *λ* ≤ 1 are parameters: *α*_1_ is the initial learning rate, *α*_2_ is the final learning rate, and *λ* is the eligibility trace [1, 17].

In the special case where *λ* = 1, the update of initial state-action values becomes

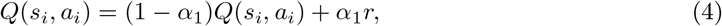

that is, the estimated values of choosing *left* and *right* in each initial state are updated independently of the final state’s estimated value. Thus, SARSA (*λ* = 1) ignores the identity of the final state when making initial-state decisions, and an initial-state action that resulted in a reward will necessarily lead to a higher stay probability when the respective initial state recurs. This is true even if the action will probably lead to the final state with the lowest value.

### 4.4 Model-based algorithm

In simulations of model-based agents [17], values were assigned to initial-state actions and to final states. The value *V* of a final state *s* ∈ {*pink, blue*} in the first trial *t* =1 was *V*(*s*, 1) = 0.5. An initial-state choice *c* ∈ {*left, right*} in trial *t* had a value *V* given by

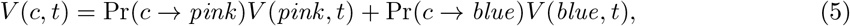

where Pr(*c* → *s*) is the probability that choosing *c* will lead to the final state *s*, which might be 0.8 or 0.2 according to the task’s transition model. The value of an initial-state choice can thus be understood as the expected value of the final state the agent will go to after making that choice. If *V*(*left, t*) > *V*(*right, t*), the agent was more likely to choose left and vice-versa.

In each trial *t*, the agent’s initial state action was *left* with probability *p_left_* and *right* with probability 1 − *p_left_*, given by

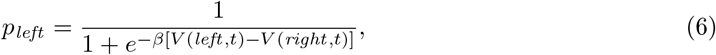

where *β* is an inverse temperature parameter. After the agent made its initial-state choice and went to a final state *s*, that final state’s value was updated according to the following equation:

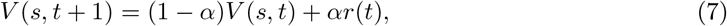

where *r*(*t*) ∈ {0, 1} indicates if the agent received a reward and 0 ≤ *α* ≤ 1 is a learning-rate parameter of the model. The value of a final state is thus the moving average of the rewards received in that state.

### 4.5 Data analysis by logistic regression

For each human participant or simulated agent, we calculated the stay probability in pairs of consecutive trials as a function of reward, transition, initial-state choice and visited final state in the first trial [34]. In the second trial of each pair, if the human participant or simulated agent chose an action (left or right) that was the same as that chosen in the previous trial, this was considered a stay. For each trial pair, the second trial’s choice was coded as the random variable *y* and classified as a stay (*y* = 1) or not a stay (*y* = 0). For each condition, trial pairs were divided into four subsets: “same letters” (if the letters presented in the first trial were the same as the letters presented in the second trial; for example, AB for the first trial, AB for the second), “same first letter” (if the first letter presented in the first trial was the same as the first letter presented in the second trial, but the second letter was different; for example, AB for the first trial, AA for the second), “same second letter” (if the second letter presented in the first trial was the same as the second letter presented in the second trial, but the first letter was different; for example, AB for the first trial, BB for the second), and “different letters” (if both letters presented in the first trial were different from the letters presented in the second trial; for example, AB for the first trial, BA for the second). For each trial pair subset, a separate analysis was performed.

We then analyzed the resulting data using a hierarchical logistic regression model whose parameters were estimated through Bayesian computational methods. The dependent variable was *p*_stay_, the stay probability for a given trial, and the independent variables were *x_r_*, which indicated whether a reward was received or not in the previous trial (+1 if the previous trial was rewarded, − 1 otherwise), *x_t_*, which indicated whether the transition in the previous trial was common or rare (+1 if it was common, −1 if it was rare), the interaction between the two, *x_c_*, which indicated whether in the previous trial the participant chose or not the initial-state choice with the highest reward probability (+1 if the choice had the highest reward probability, −1 otherwise), and *x_f_*, which indicated whether in the pervious trial the participant visited the final state with the highest reward probability (+1 if the final state had the highest reward probability, −1 otherwise). Thus, for each condition, we determined a intercept 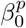 for each participant and five fixed coefficients that are shown in the following equation:

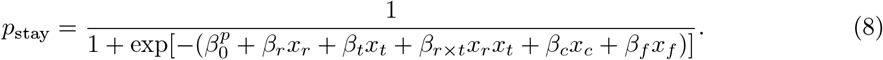

The distribution of *y* was Bernoulli(*p*_stay_). The distribution of the 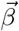 vectors was 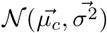 if the participant was in the simultaneous condition and 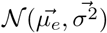 if the participant was in the sequential condition; in other words, the subset means for each 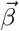 were allowed to vary independently. The parameters of the 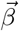 distribution were given vague prior distributions based on preliminary analyses—the 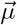 vectors’ components were given a 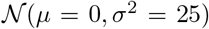 prior, and the 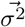 vector’s components were given a Half-normal(0, 25) prior. Other vague prior distributions for the model parameters were tested and the results did not change significantly.

To obtain parameter estimates from the model’s posterior distribution, we coded the model into the Stan modeling language [41, 42] and used the PyStan Python package [43] to obtain 80,000 samples of the joint posterior distribution from four chains of length 40,000 (warmup 20,000). Convergence of the chains was indicated by 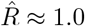 for all parameters.

### 4.6 Fitting of the algorithms to experimental data

For comparison with the participant data, we fitted the SARSA model-free algorithm and the model-based algorithm to the experimental data and generated replicated data using the fitted parameters. The parameters were obtained by fitting both algorithms to all participants. To that end, we used a Bayesian hierarchical model, which allowed us to pool data from all participants to improve individual parameter estimates.

The parameters of the model-based algorithm for the *ith* participant were *α^i^* and *β^i^*. They were given a Beta(*a_α_, b_α_*) and 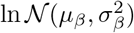 prior distributions respectively. The hyperparameters *a_α_* and *b_α_* were themselves given a noninformative Half-normal(0, 10^4^) prior and the hyperparameters *μ_β_* and 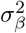 were given a noninformative 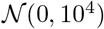 and Half-normal(0, 10^4^) priors respectively. The parameters of the model-free algorithm for the *i*th participant were 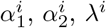, and *β^i^*. They were given a Beta(*a*_*α*_1__, *b*_*α*_1__), Beta(*a*_*α*_2__, *b*_*α*_1__), Beta(*a_λ_, b_λ_*) and 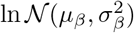 prior distributions respectively. The hyperparameters *a*_*α*_1__, *a*_*α*_2__, *a_λ_, b*_*α*_1__, *b*_*α*_2__, and *b_λ_* were themselves given a noninformative Half-normal(0, 10^4^) prior and the hyperparameters *μ_β_* and 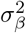 were given a noninformative 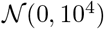 and Half-normal(0, 10^4^) priors respectively. We then coded the models into the Stan modeling language [41, 42] and used the PyStan Python package [43] to obtain 40,000 samples of the joint posterior distribution from one chain of length 80,000 (warmup 40,000). Convergence of the chains was indicated by 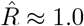 for all parameters.

### 4.7 Code and data availability

All the behavioral data used in this study are available at https://github.com/carolfs/mf_wm

## Appendix

**Figure 11:**
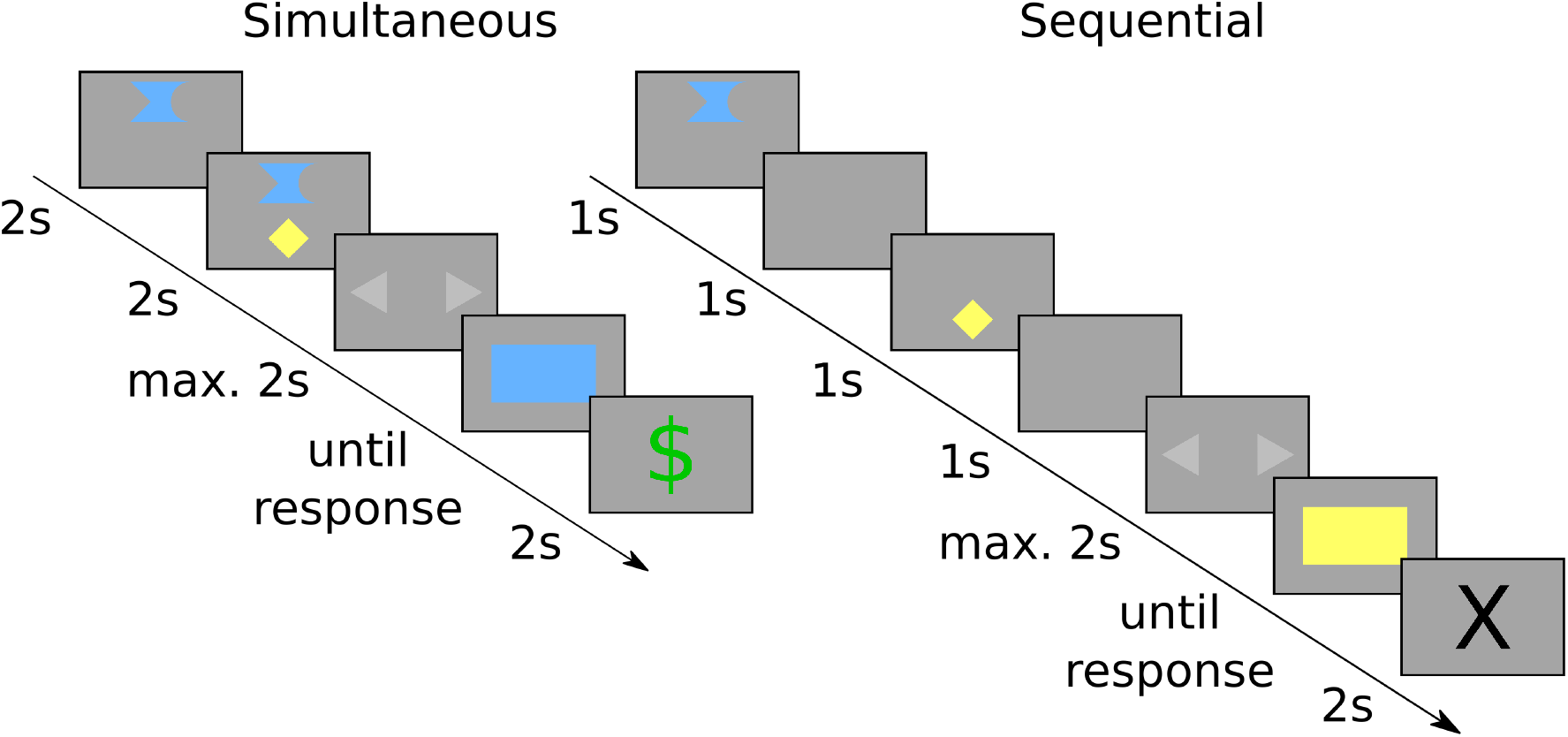
Timeline of a replication of the current study using figures to identify the initial states rather than letters. The two final states were the blue state and the yellow state.

**Figure 12:**
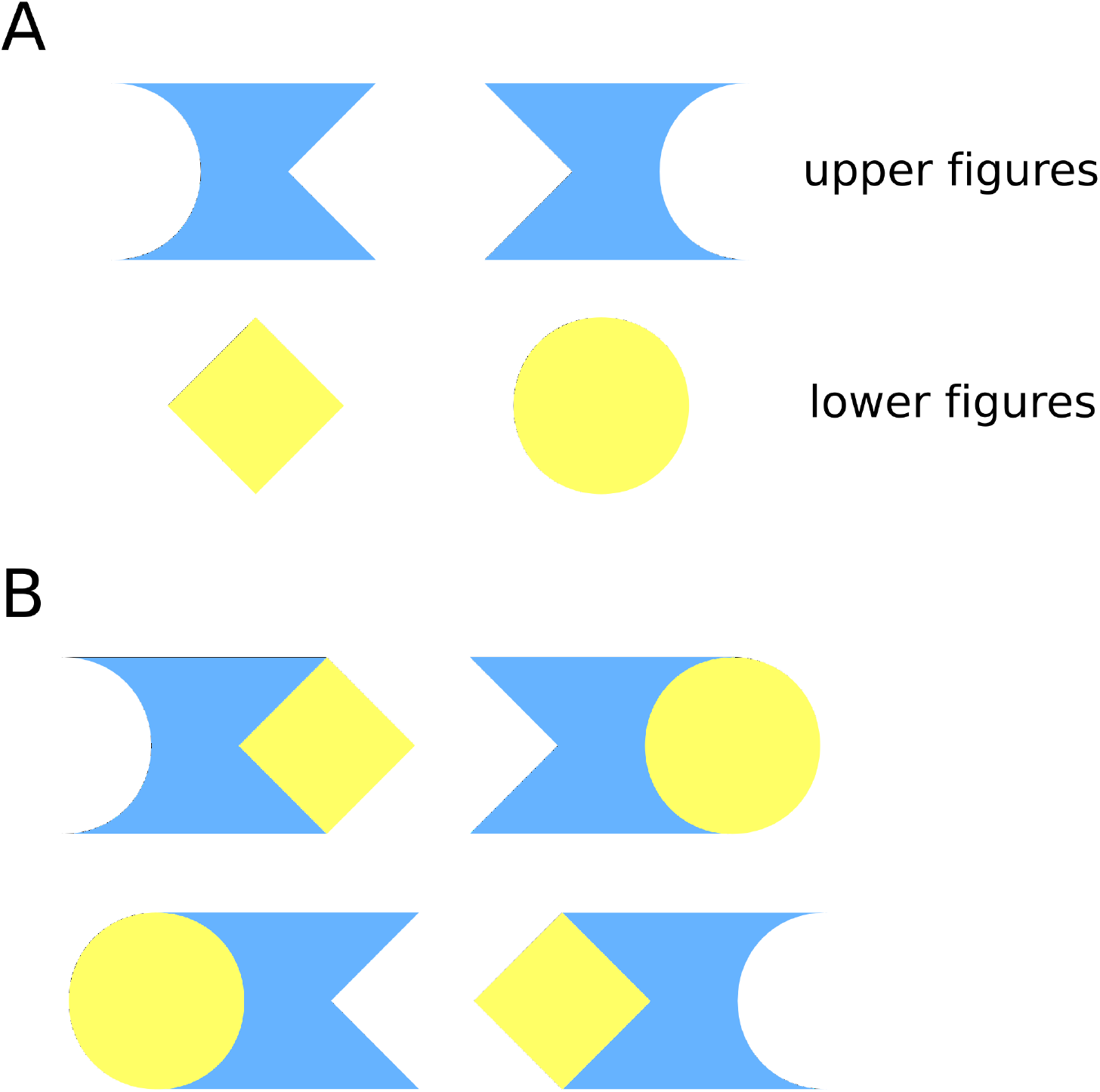
In our replication task, participants saw two figures on the screen, an upper figure and a lower figure (A). They were instructed to imagine how the two figures would fit together in other to determine the common and rare transitions (B). If, for instance, the participant saw one of the upper pair of figures shown in panel B, pressing left would commonly take them to the blue final state and pressing right would commonly take them to the yellow final state. If instead they saw one of the lower pair of figures in panel B, pressing right would commonly take them to yellow state and vice versa.

**Figure 13:**
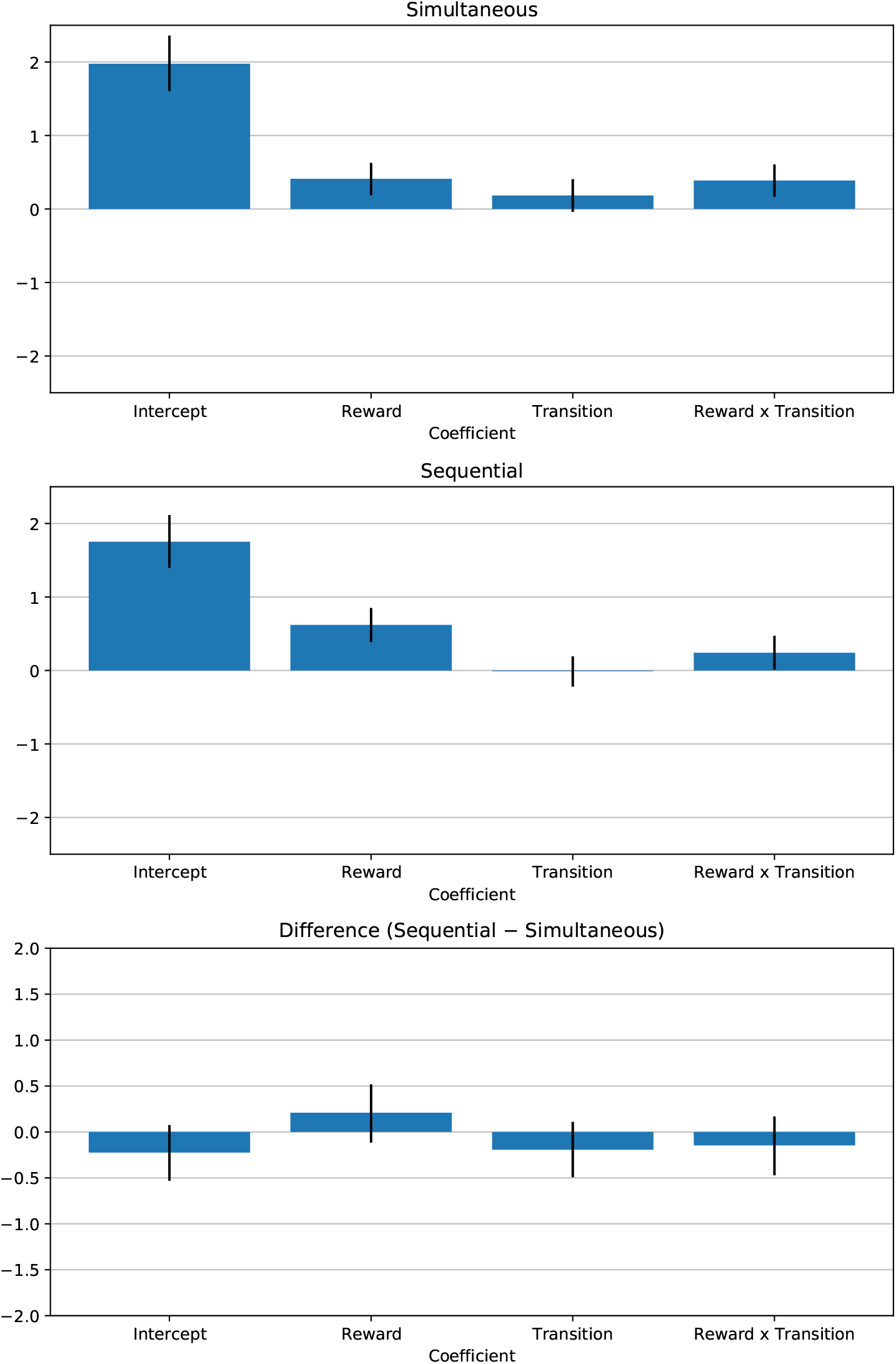
Logistic regression coefficients of human participants in our replication experiment for consecutive trial pairs in the “same letters” subset. The error bars correspond to the 95% credible interval.

**Figure 14:**
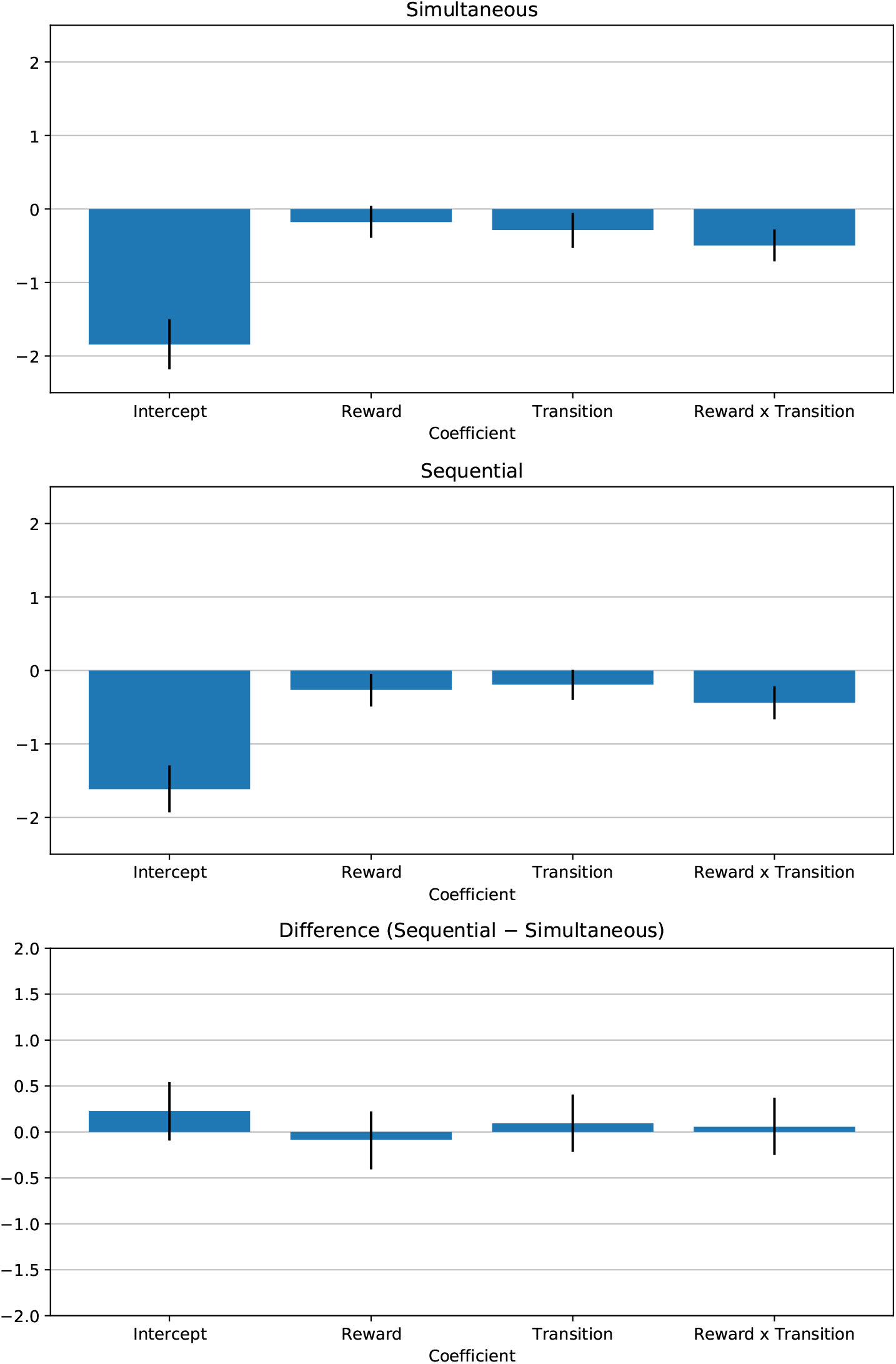
Logistic regression coefficients of human participants in our replication experiment for consecutive trial pairs in the “same first letter” subset. The error bars correspond to the 95% credible interval.

**Figure 15:**
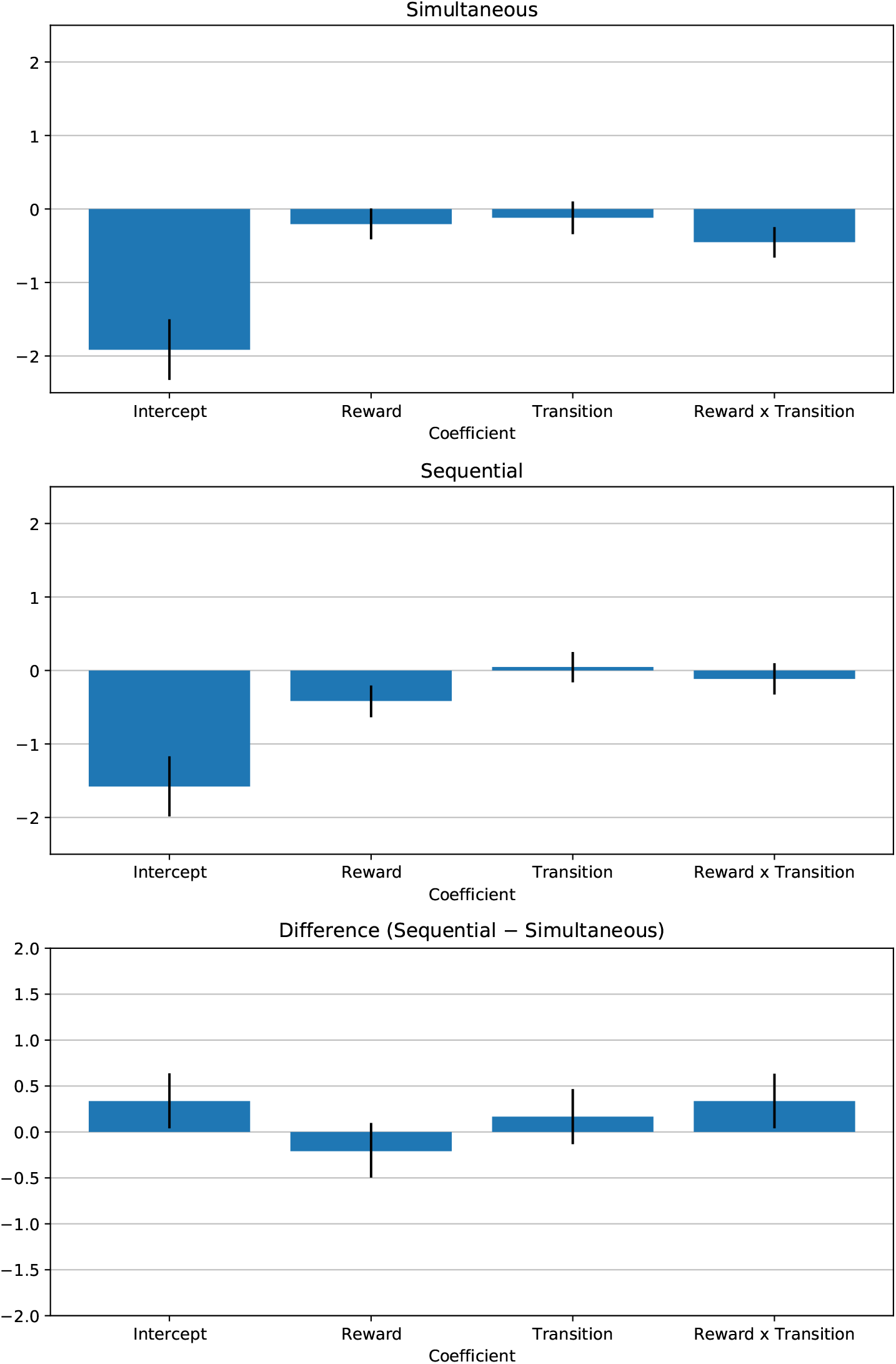
Logistic regression coefficients of human participants in our replication experiment for consecutive trial pairs in the “same second letter” subset. The error bars correspond to the 95% credible interval.

**Figure 16:**
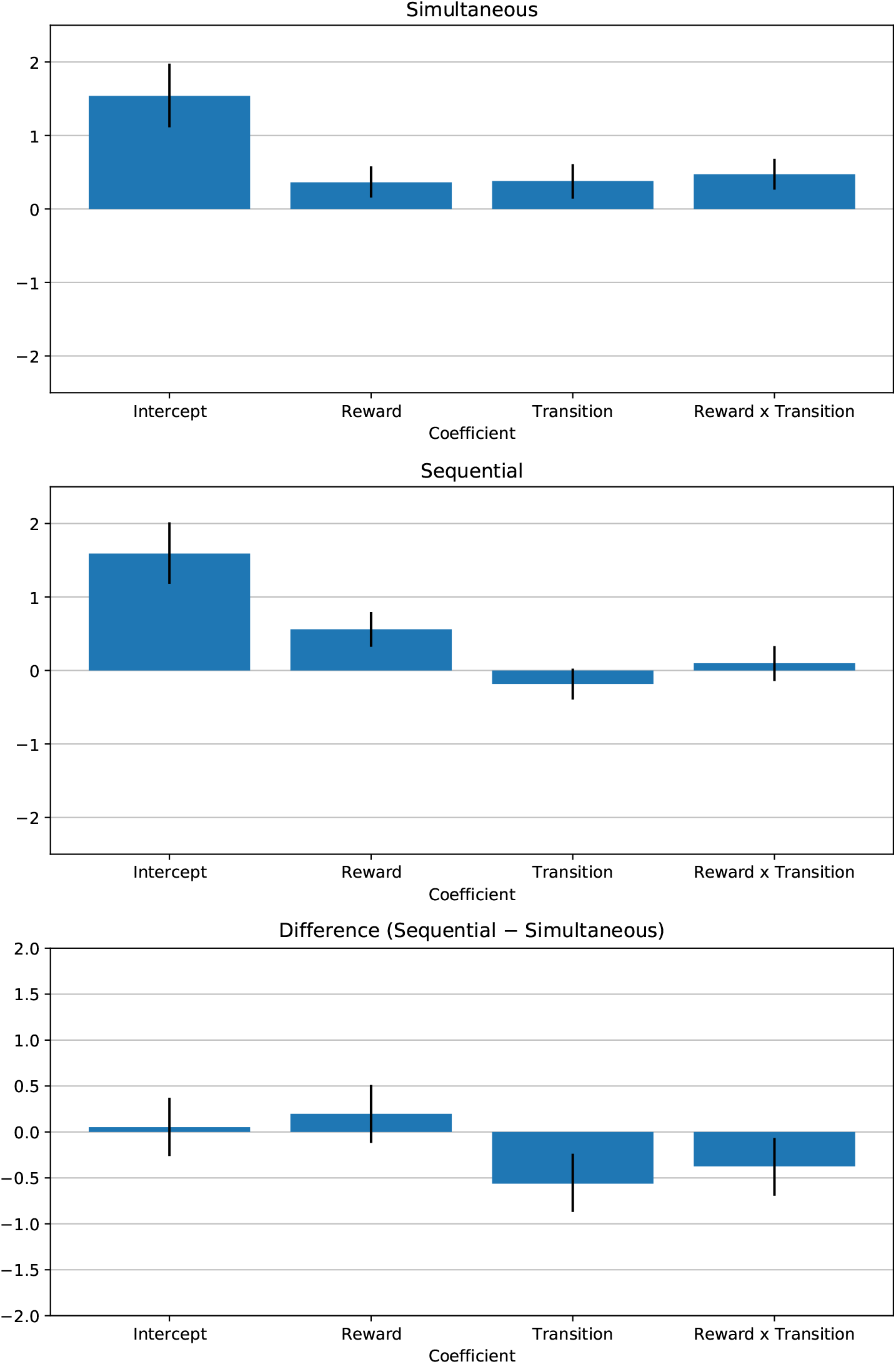
Logistic regression coefficients of human participants in our replication experiment for consecutive trial pairs in the “different letters” subset. The error bars correspond to the 95% credible interval.

## References

[1] Richard S. Sutton and Andrew G. Barto. Reinforcement Learning: An Introduction. A Bradford Book, first edition, 1998.

[2] W. Schultz, P. Dayan, and P. R. Montague. A Neural Substrate of Prediction and Reward. Science, 275(5306):1593–1599, mar 1997. ISSN 0036-8075. doi: 10.1126/science.275.5306.1593. URL http://www.sciencemag.org/cgi/doi/10.1126/science.275.5306.1593.

[3] Christopher D Fiorillo, Philippe N Tobler, and Wolfram Schultz. Discrete coding of reward probability and uncertainty by dopamine neurons. Science (New York, N.Y.), 299(5614):1898–902, mar 2003. ISSN 1095-9203. doi: 10.1126/science.1077349. URL http://www.ncbi.nlm.nih.gov/pubmed/12649484.

[4] Yael Niv. Reinforcement learning in the brain. Journal of Mathematical Psychology, 53(3):139–154, jun 2009. ISSN 00222496. doi: 10.1016/j.jmp.2008.12.005. URL http://linkinghub.elsevier.com/retrieve/pii/S0022249608001181.

[5] Paul W. Glimcher. Understanding dopamine and reinforcement learning: The dopamine reward prediction error hypothesis. Proceedings of the National Academy of Sciences, 108 (Supplement_3):15647–15654, sep 2011. ISSN 0027-8424. doi: 10.1073/pnas.1014269108. URL http://www.pnas.org/cgi/doi/10.1073/pnas.1014269108.

[6] Daeyeol Lee, Hyojung Seo, and Min Whan Jung. Neural Basis of Reinforcement Learning and Decision Making. Annual Review of Neuroscience, 35(1):287–308, jul 2012. ISSN 0147-006X. doi: 10.1146/annurev-neuro-062111-150512. URL http://www.annualreviews.org/doi/abs/10.1146/annurev-neuro-062111-150512.

[7] Ray J. Dolan and Peter Dayan. Goals and Habits in the Brain. Neuron, 80(2):312–325, oct 2013. ISSN 08966273. doi: 10.1016/j.neuron.2013.09.007. URL http://linkinghub.elsevier.com/retrieve/pii/S0896627313008052.

[8] Ann M. Graybiel. Habits, Rituals, and the Evaluative Brain. Annual Review of Neuroscience, 31(1):359–387, jul 2008. ISSN 0147-006X. doi: 10.1146/annurev.neuro.29.051605.112851. URL http://www.annualreviews.org/doi/10.1146/annurev.neuro.29.051605.112851.

[9] Peter Dayan. How to set the switches on this thing. Current Opinion in Neurobiology, 22(6):1068–1074, dec 2012. ISSN 09594388. doi: 10.1016/j.conb.2012.05.011. URL http://linkinghub.elsevier.com/retrieve/pii/S0959438812000992.

[10] Kyle S. Smith and Ann M. Graybiel. Investigating habits: strategies, technologies and models. Frontiers in Behavioral Neuroscience, 8, 2014. ISSN 1662-5153. doi: 10.3389/fnbeh.2014. 00039. URL http://journal.frontiersin.org/article/10.3389/fnbeh.2014.00039/ abstract.

[11] Fiery Cushman and Adam Morris. Habitual control of goal selection in humans. Proceedings of the National Academy of Sciences, 112(45):13817–13822, nov 2015. ISSN 0027-8424. doi: 10.1073/pnas.1506367112. URL http://www.pnas.org/lookup/doi/10.1073/pnas.1506367112.

[12] Henk Aarts and Ap Dijksterhuis. Habits as knowledge structures: Automaticity in goal-directed behavior. Journal of Personality and Social Psychology, 78(1):53–63, 2000. ISSN 1939-1315. doi: 10.1037/0022-3514.78.1.53. URL http://doi.apa.org/getdoi.cfm?doi=10.1037/0022-3514.78.1.53.

[13] Amir Dezfouli and Bernard W. Balleine. Habits, action sequences and reinforcement learning. European Journal of Neuroscience, 35(7):1036–1051, apr 2012. ISSN 0953816X. doi: 10. 1111/j.1460-9568.2012.08050.x. URL http://doi.wiley.com/10.1111/j.1460-9568.2012.08050.x.

[14] Amir Dezfouli and Bernard W. Balleine. Actions, Action Sequences and Habits: Evidence That Goal-Directed and Habitual Action Control Are Hierarchically Organized. PLoS Computational Biology, 9(12):e1003364, dec 2013. ISSN 1553-7358. doi: 10.1371/journal.pcbi.1003364. URL http://dx.plos.org/10.1371/journal.pcbi.1003364.

[15] A. Dezfouli, N. W. Lingawi, and B. W. Balleine. Habits as action sequences: hierarchical action control and changes in outcome value. Philosophical Transactions of the Royal Society B: Biological Sciences, 369(1655):20130482–20130482, sep 2014. ISSN 0962-8436. doi: 10. 1098/rstb.2013.0482. URL http://rstb.royalsocietypublishing.org/cgi/doi/10.1098/rstb.2013.0482.

[16] Randall C. O’Reilly and Michael J. Frank. Making Working Memory Work: A Computational Model of Learning in the Prefrontal Cortex and Basal Ganglia. Neural Computation, 18 (2):283–328, feb 2006. ISSN 0899-7667. doi: 10.1162/089976606775093909. URL http://www.mitpressjournals.org/doi/abs/10.1162/089976606775093909.

[17] Nathaniel D. Daw, Samuel J. Gershman, Ben Seymour, Peter Dayan, and Raymond J. Dolan. Model-Based Influences on Humans’ Choices and Striatal Prediction Errors. Neuron, 69(6): 1204–1215, mar 2011. ISSN 08966273. doi: 10.1016/j.neuron.2011.02.027. URL http://linkinghub.elsevier.com/retrieve/pii/S0896627311001255.

[18] Michael T Todd, Yael Niv, and Jonathan D Cohen. Learning to Use Working Memory in Partially Observable Environments through Dopaminergic Reinforcement. In D Koller, D Schuurmans, Y Bengio, and L Bottou, editors, Advances in Neural Information Processing Systems 21, pages 1689–1696. Curran Associates, Inc., 2009. URL http://papers.nips.cc/paper/3508-learning-to-use-working-memory-in-partially-observable-environments-through-dopaminergic-reinforcement.pdf.

[19] Arthur S. Reber. Implicit learning and tacit knowledge. Journal of Experimental Psychology: General, 118(3):219–235, 1989. ISSN 1939-2222. doi: 10.1037/0096-3445.118.3.219. URL http://doi.apa.org/getdoi.cfm?doi=10.1037/0096-3445.118.3.219.

[20] Richard L. Canfield and Marshall M. Haith. Young infants’ visual expectations for symmetric and asymmetric stimulus sequences. Developmental Psychology, 27(2):198–208, 1991. ISSN 0012-1649. doi: 10.1037/0012-1649.27.2.198. URL http://doi.apa.org/getdoi.cfm?doi=10.1037/0012-1649.27.2.198.

[21] Scott A. Huettel, Peter B. Mack, and Gregory McCarthy. Perceiving patterns in random series: dynamic processing of sequence in prefrontal cortex. Nature Neuroscience, apr 2002. ISSN 10976256. doi: 10.1038/nn841. URL http://www.nature.com/doifinder/10.1038/nn841.

[22] Asher Cohen, Richard I. Ivry, and Steven W. Keele. Attention and structure in sequence learning. Journal of Experimental Psychology: Learning, Memory, and Cognition, 16(1):17–30, 1990. ISSN 1939-1285. doi: 10.1037/0278-7393.16.1.17. URL http://doi.apa.org/getdoi.cfm?doi=10.1037/0278-7393.16.1.17.

[23] Axel Cleeremans and James L. McClelland. Learning the structure of event sequences. Journal of Experimental Psychology: General, 120(3):235–253, 1991. ISSN 1939-2222. doi: 10.1037/0096-3445.120.3.235. URL http://doi.apa.org/getdoi.cfm?doi=10.1037/0096-3445.120.3.235.

[24] I H Jenkins, D J Brooks, P D Nixon, R S Frackowiak, and R E Passingham. Motor sequence learning: a study with positron emission tomography. The Journal of neuroscience : the official journal of the Society for Neuroscience, 14(6):3775–90, jun 1994. ISSN 0270-6474. URL http://www.ncbi.nlm.nih.gov/pubmed/8207487.

[25] Eli Vakil, Shimon Kahan, Moshe Huberman, and Alicia Osimani. Motor and non-motor sequence learning in patients with basal ganglia lesions: The case of serial reaction time (SRT). Neuropsychologia, 38(1):1–10, 2000. ISSN 00283932. doi: 10.1016/S0028-3932(99)00058-5.

[26] S. Lehericy, H. Benali, P.-F. Van de Moortele, M. Pelegrini-Issac, T. Waechter, K. Ugurbil, and J. Doyon. Distinct basal ganglia territories are engaged in early and advanced motor sequence learning. Proceedings of the National Academy of Sciences, 102(35):12566–12571, aug 2005. ISSN 0027-8424. doi: 10.1073/pnas.0502762102. URL http://www.pnas.org/cgi/doi/10.1073/pnas.0502762102.

[27] A. R. Otto, C. M. Raio, A. Chiang, E. A. Phelps, and N. D. Daw. Working-memory capacity protects model-based learning from stress. Proceedings of the National Academy of Sciences, 110(52):20941–20946, dec 2013. ISSN 0027-8424. doi: 10.1073/pnas.1312011110. URL http://www.pnas.org/cgi/doi/10.1073/pnas.1312011110.

[28] A. Ross Otto, Samuel J. Gershman, Arthur B. Markman, and Nathaniel D. Daw. The Curse of Planning. Psychological Science, 24(5):751–761, may 2013. ISSN 0956-7976. doi: 10.1177/ 0956797612463080. URL http://journals.sagepub.com/doi/10.1177/0956797612463080.

[29] A. Ross Otto, Anya Skatova, Seth Madlon-Kay, and Nathaniel D. Daw. Cognitive Control Predicts Use of Model-based Reinforcement Learning. Journal of Cognitive Neuroscience, 27(2):319–333, feb 2015. ISSN 0898-929X. doi: 10.1162/jocn_a_00709. URL http://www.mitpressjournals.org/doi/abs/10.1162/jocn{_}a{_}00709http://www.mitpressjournals.org/doi/10.1162/jocn{_}a{_}00709.

[30] J. H. Decker, A. R. Otto, N. D. Daw, and C. A. Hartley. From Creatures of Habit to Goal-Directed Learners: Tracking the Developmental Emergence of Model-Based Reinforcement Learning. Psychological Science, 27(6):848–858, jun 2016. ISSN 0956-7976. doi: 10.1177/0956797616639301. URL http://pss.sagepub.com/lookup/doi/10.1177/0956797616639301.

[31] Peter Smittenaar, Thomas H.B. FitzGerald, Vincenzo Romei, Nicholas D. Wright, and Raymond J. Dolan.Disruption of Dorsolateral Prefrontal Cortex Decreases Model-Based in Favor of Model-free Control in Humans. Neuron, 80(4): 914–919, nov 2013. ISSN 08966273. doi: 10.1016/j.neuron.2013.08.009. URL http://linkinghub.elsevier.com/retrieve/pii/S0896627313007204.

[32] Thomas Akam, Rui Costa, and Peter Dayan. Simple Plans or Sophisticated Habits? State, Transition and Learning Interactions in the Two-Step Task. PLOS Computational Biology, 11(12): e1004648, dec 2015. ISSN 1553-7358. doi: 10.1371/journal.pcbi.1004648. URL http://dx.plos.org/10.1371/journal.pcbi.1004648.

[33] Kevin J Miller, Carlos D Brody, and Matthew M Botvinick. Identifying Model-Based and Model-Free Patterns in Behavior on Multi-Step Tasks. bioRxiv, page 14, 2016. doi: 10.1101/ 096339. URL https://doi.org/10.1101/096339.

[34] Carolina Feher da Silva and Todd A. Hare. A note on the analysis of two-stage task results: How changes in task structure affect what model-free and model-based strategies predict about the effects of reward and transition on the stay probability. PLOS ONE, 13(4):e0195328, apr 2018. ISSN 1932-6203. doi: 10.1371/journal.pone.0195328. URL http://dx.plos.org/10.1371/journal.pone.0195328.

[35] Bradley B Doll, Katherine D Duncan, Dylan A Simon, Daphna Shohamy, and Nathaniel D Daw. Model-based choices involve prospective neural activity. Nature Neuroscience, 18(5): 767–772, mar 2015. ISSN 1097-6256. doi: 10.1038/nn.3981. URL http://www.nature.com/doifinder/10.1038/nn.3981.

[36] Wouter Kool, Fiery A. Cushman, and Samuel J. Gershman. When Does Model-Based Control Pay Off? PLOS Computational Biology, 12(8):e1005090, aug 2016. ISSN 1553-7358. doi: 10.1371/journal.pcbi.1005090. URL http://dx.plos.org/10.1371/journal.pcbi.1005090.

[37] G. Cumming. Precision for Planning. In Understanding The New Statistics, chapter 13, pages 355–380. Routledge, New York, London, 1 edition, 2012.

[38] J. K. Kruschke. Goals, Power, and Sample Size. In Doing Bayesian Data Analysis, chapter 13, pages 359–398. Academic Press, London, 2 edition, 2015.

[39] Nathaniel D Daw, Yael Niv, and Peter Dayan. Uncertainty-based competition between pre-frontal and dorsolateral striatal systems for behavioral control. Nature neuroscience, 8(12): 1704–11, dec 2005. ISSN 1097-6256. doi: 10.1038/nn1560. URL http://dx.doi.org/10.1038/nn1560http://www.ncbi.nlm.nih.gov/pubmed/16286932.

[40] Nathaniel D. Daw and John P. O’Doherty.Multiple Systems for Value Learning. In Paul W. Glimcher and Ernst Fehr, editors, Neuroeconomics, chapter 21, pages 393–410. Elsevier, second edition, 2014. doi: 10.1016/B978-0-12-416008-8.00021-8. URL http://linkinghub.elsevier.com/retrieve/pii/B9780124160088000218.

[41] Bob Carpenter, Andrew Gelman, Matthew D. Hoffman, Daniel Lee, Ben Goodrich, Michael Betancourt, Marcus Brubaker, Jiqiang Guo, Peter Li, and Allen Riddell. Stan: A Probabilistic Programming Language. Journal of Statistical Software, 76(1), 2017. ISSN 1548-7660. doi: 10.18637/jss.v076.i01. URL http://www.jstatsoft.org/v76/i01/.

[42] Stan Development Team. Stan Modeling Language Users Guide and Reference Manual, Version 2.16.0, 2017.

[43] Stan Development Team. PyStan: the Python interface to Stan, 2017. URL http://mc-stan.org.

